# Onset of differentiation is posttranscriptionally controlled in adult neural stem cells

**DOI:** 10.1101/289868

**Authors:** Avni Baser, Yonglong Dang, Maxim Skabkin, Gülce S. Gülcüler Balta, Georgios Kalamakis, Susanne Kleber, Manuel Göpferich, Roman Schefzik, Alejandro Santos Lopez, Enric Llorens Bobadilla, Carsten Schultz, Bernd Fischer, Ana Martin-Villalba

## Abstract

The contribution of posttranscriptional regulation of gene expression to neural stem cell differentiation during tissue homeostasis remains elusive. Here we show highly dynamic changes in protein synthesis along differentiation of stem cells to neurons *in vivo*. Examination of individual transcripts using RiboTag mouse models reveals that neural stem cells efficiently translate abundant transcripts, whereas translation becomes increasingly controlled with the onset of differentiation. Stem cell generation of early neuroblasts involves translational repression of a subset of mRNAs including the stem cell-identity factors Sox2 and Pax6 as well as translation machinery components. *In silico* motif analysis identifies a pyrimidine-rich motif (PRM) in this repressed subset. A drop in mTORC1 activity at the onset of differentiation selectively blocks translation of PRM-containing transcripts. Our data uncovers how a drop in mTORC1 allows robust simultaneous posttranscriptional repression of key stem cell identity-factors and translation-components and thereby stemness exit and migration.

---

To generate olfactory bulb interneurons in the adult brain, ventricle-contacting NSCs exit the dormant state to transit through a primed quiescent to an active state^1–3^ and produce neurogenic progenitors (NPs). NPs differentiate into neuroblasts (NBs) that migrate along the rostral migratory stream to the olfactory bulb (OB) where they ultimately transform into new granule and periglomerular interneurons^4,5^. Maintenance of NSCs throughout the life of the animal critically depends on keeping them in a state of quiescence, which they entered during embryonic development^6,7^. This dormant state features low protein synthesis^2^ that is key to maintain the pool of fully functional stem cells not only in the brain but also in the bone marrow (haematopoietic stem cells: HSCs) and hair follicles (hair follicle stem cells: HFSCs)^8–10^. With transition into the primed state, protein synthesis increases before actual division^2,8,11^. However, how protein synthesis changes along the further steps of maturation has not been studied.

## Global protein synthesis is highly dynamic along stem cell differentiation to neurons

To address active protein synthesis during adult olfactory bulb neurogenesis, we FACS-isolated multiple cell populations from the ventricular-subventricular zone (SVZ) and the olfactory bulb based on their surface marker expression as previously described^2^. This strategy enables distinction between quiescent NSCs (qNSCs, GLAST^+^PROM^+^), active NSCs (aNSCs, GLAST^+^PROM^+^EGFR^+^), neurogenic progenitors (NPs, only EGFR^+^), early neuroblasts (ENBs, PSA-NCAM^+^ from SVZ) and late neuroblasts (LNBs, PSA-NCAM^+^ from OB, Fig. 1a). In addition, to overcome the challenging need of sorting neurons we isolated cells from the olfactory bulb and identified by marker expression post fixation: LNBs (DCX^+^NeuN^−^), early neurons (DCX^+^NeuN^+^) and mature neurons (DCX^−^NeuN^+^) (Fig. 1d). Active protein synthesis per cell was quantified by incorporation of the aa-tRNA mimic homologue O-Propargyl-Puromycin (OPP) into nascent proteins^12^. We confirmed that global protein synthesis is very low in qNSCs and increases dramatically in aNSCs^2^ (Fig. 1b-c). In addition, we identified a further increase in neurogenic progenitors and a dramatic drop in ENBs back to levels found in qNSC (Fig. 1b-c). To directly assess global protein synthesis in ENB that exclusively derive from NSCs we used Tlx-CreER-eYFP reporter mice. Tlx-recombined population overlapped with the previously characterized GLAST+PROM+ population of NSC, confirming that we are looking at the same subset of cells with Tlx-labeling and FACS-isolation approach (Extended data Fig. 1a,b). Reporter-positive ENBs exhibited a dramatic drop in OPP incorporation as compared to NSCs, similar to the drop found in FACS-isolated populations (Extended data Fig. 1c,d). Thereafter, active protein synthesis increases again in LNBs, which in contrast to ENBs are postmitotic and, thus, we hypothesize that this increase potentially relates to the complex process of integration into the neuronal network. Altogether these data support the idea that the need to differentiate and not proliferation dictates the level of protein synthesis as previously reported for HFSCs^8^. From LNBs to early and mature neurons active protein synthesis gradually decreases (Fig. 1 e,f), supposedly because neurons rather need local proteome changes through localized translation^13^. Indeed, a recent study addressing the axonal translatome by ribotagging demonstrates that a subset of mRNAs with key axon-specific functions is highly enriched in axons where they are locally translated^14^.

**Figure 1.**
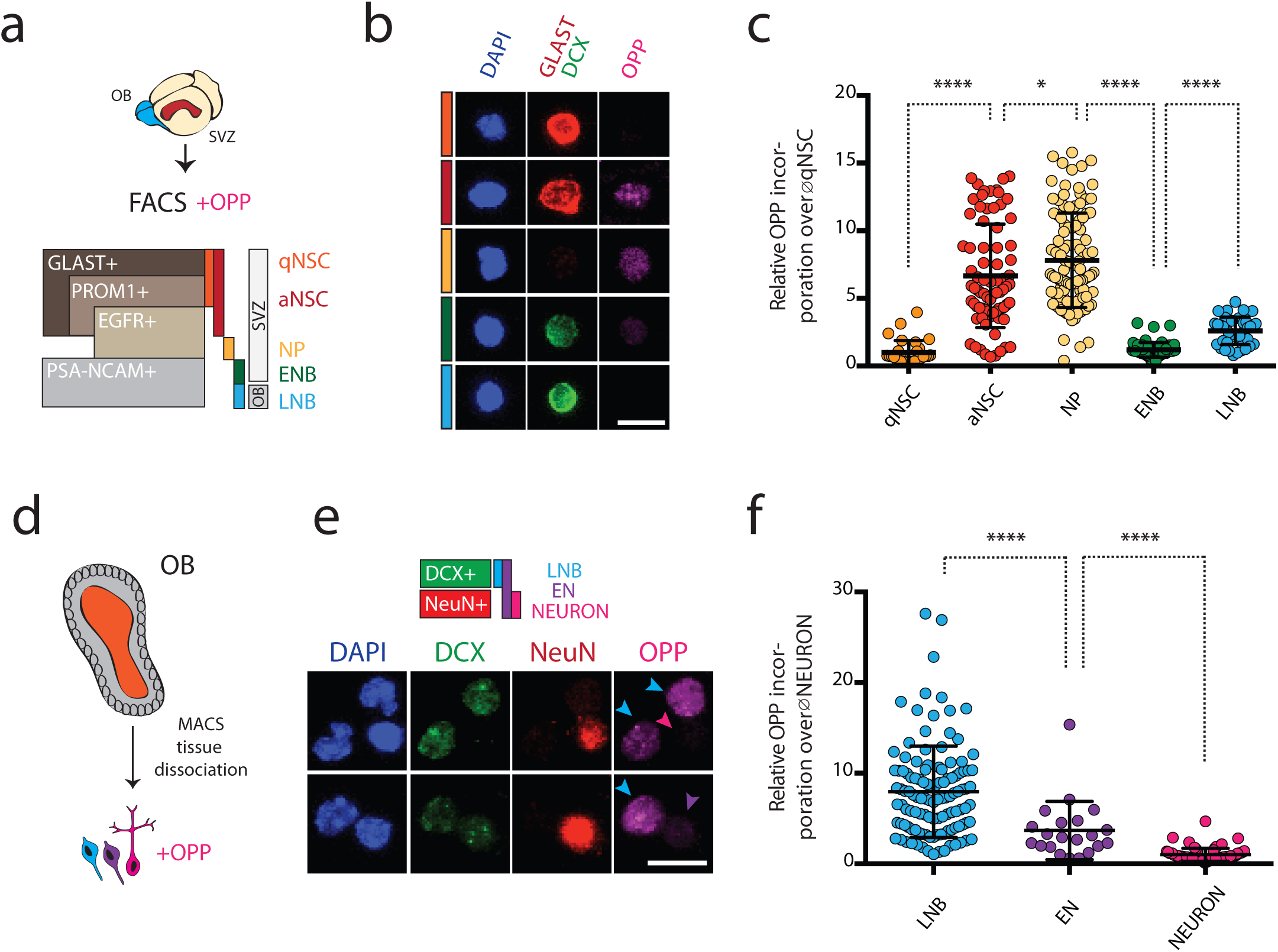
Global protein synthesis is highly dynamic during neurogenesis. **a,** Sorting strategy for different stages of NSC differentiation. **b,** Representative images of OPP incorporation with specific marker expression, GLAST for NSCs, DCX for neuroblasts. **c,** Quantification of OPP incorporation relative to qNSCs. **d,** Strategy to compare protein synthesis of olfactory bulb neuroblasts (LNB; DCX^+^NeuN^−^) to early neurons (EN; DCX^+^NeuN^+^) and mature neurons (DCX^−^NeuN^+^). **e,** Representative images of OPP incorporation of LNBs, ENs and mature neurons. **f,** Quantification of OPP incorporation of LNBs, ENs and neurons (relative to neurons). qNSC: quiescent NSC, aNSC: active NSC, NP: neurogenic progenitor, ENB: early neuroblast (SVZ), LNB: late neuroblast (OB), EN: early neuron, OPP: O-Propargyl-puromycin. Statistical significance by student’s t-test (Mann-Whitney). All scale bars: 10µm.

## RiboTag mouse models target distinct stages of neuronal differentiation

To address how transcript-specific translation relates to transcript abundance, we developed a system for parallel assessment of the transcriptome and translatome of NSCs and their progeny *in situ*. We used the RiboTag mouse line that carries a hemagglutinin (HA)-tagged variant of the large ribosomal subunit RPL22 (RPL22-HA)^15^, one of the most stable ribosomal components^16^ that is localized on the exposed solvent side of 60S and is not involved in either mRNA or tRNA binding or ribosomal dynamics during translation^17^. Ribotag mice were bred with previously established cre-inducible fluorescent reporter mouse lines, allows parallel ribosomal tagging and FACS isolation of the cell population of interest. We generated Tlx-CreER-Rpl22.HA-eYFP (TiCRY) and Dcx-CreER-Rpl22.HA-eYFP (DiCRY) mouse lines to target NSCs and their progeny, respectively (Fig. 2a).

**Figure 2.**
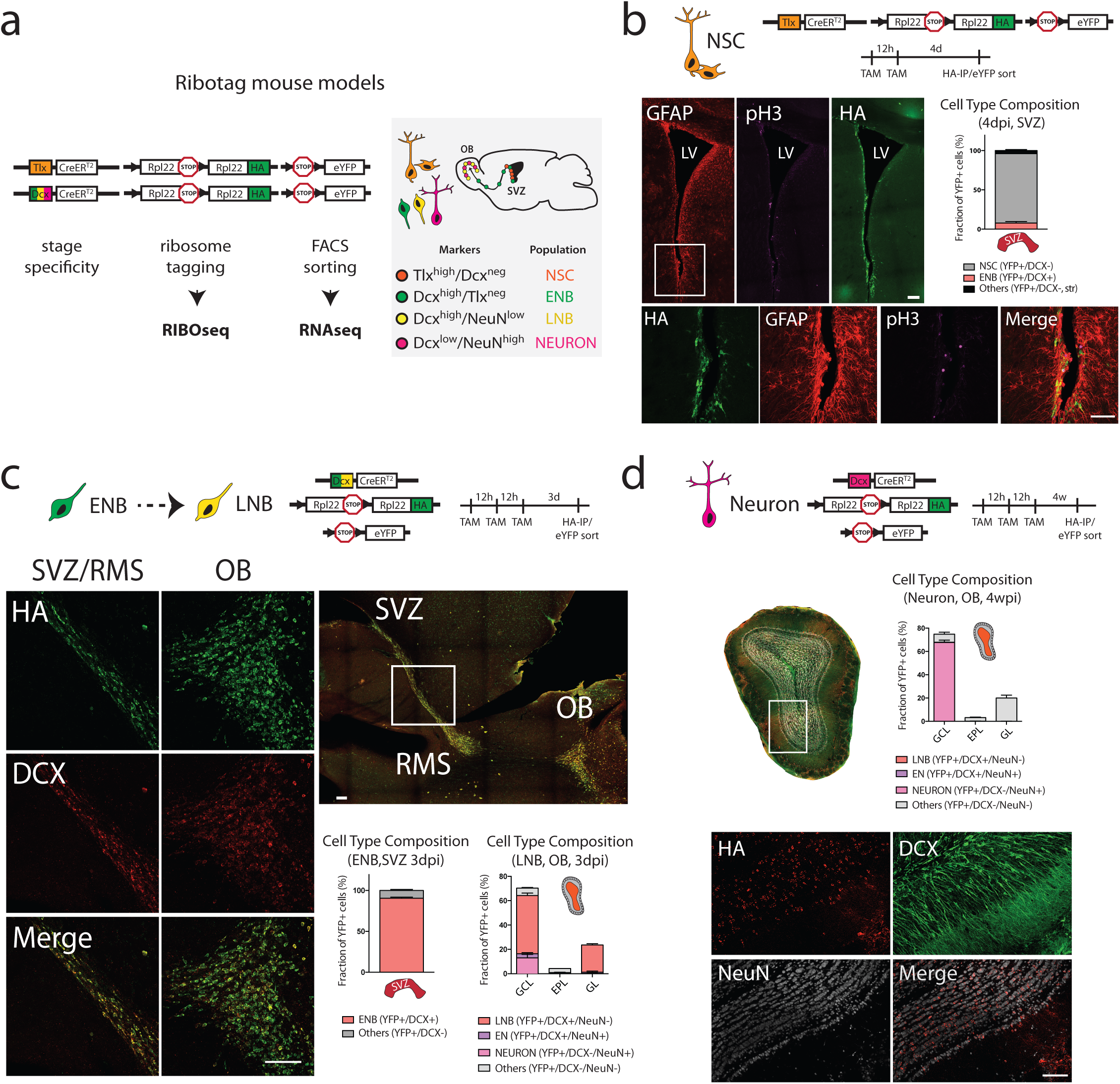
RiboTag mouse models target stages of neuronal differentiation. **a,** Schematic representation of RiboTag mouse lines used for the study, HA-tag allows isolation of ribosome-associated mRNA, cell sorting based on eYFP expression alllows isolation of total mRNA. **b,** IHC confirming HA-tag expression in NSCs (controlled by the Tlx-promotor) of the SVZ, labeled cells are mostly GFAP^+^pH3^−^. Quantification shows that labeled cells in the TiCRY-SVZ are mostly NSCs (eYFP^+^/DCX^−^: 88.32%±1.8), some ENBs (eYFP^+^/DCX^+^: 7.75%±1.9) and few of unknown identity further away from the ventricle (eYFP^+^/DCX^−^: 3.94%±1.3) at the timepoint of the experiments (4dpi). n=3. Scale bars: 50µm. **c,** IHC confirming expression of HA-tag in migrating neuroblasts (controlled by the Dcx-promotor) both in the SVZ/RMS (ENB) as well as in the OB (LNB). Quantification shows that labeled cells within the DiCRY-SVZ are mostly ENBs (eYFP^+^/DCX^+^: 90.52%±1.3) and some DCX negative cells of unknown identity (eYFP^+^/DCX^−^: 9.49%±1.3). DiCRY-olfactory bulb labeled-cells could be detected in all layers (GCL/MCL: 70.33%±1.3, EPL: 4.25%±0.3, PGL: 23,60%±1) indicating no labeling bias. eYFP cells in the GCL/MCL were mostly LNBs (eYFP^+^/DCX^+^: 47.83%±2), few early neurons (eYFP^+^/DCX^+^/NeuN^+^: 3.4%±0.9), some neurons (eYFP^+^/DCX^−^/NeuN^+^: 13%±2.9) and some cells of unknown identity (eYFP^+^/DCX^−^/NeuN^−^: 6.1%±0.5), which most likey consisted of NeuN-negative subtypes of neurons as assessed by morphology and position. Few labeled-cells could be found in the EPL. These were mostly NeuN-negative neurons (eYFP^+^/DCX^−^/NeuN^−^: 3.33%±0.1), few LNBs (eYFP^+^/DCX^+^/NeuN^−^: 0.21%±0.1) and rarely NeuN-positive neurons (eYFP^+^/DCX^−^/NeuN^+^: 0.03%±0.06). YFP-cells in the PGL were mostly LNBs (eYFP^+^/DCX^+^/NeuN^−^: 22.4%±1), some early neurons (eYFP^+^/DCX^+^/NeuN^+^: 1%±0.9) and rarely mature neurons (eYFP^+^/DCX^−^/NeuN^+^: 0.2%±0.2). n=3. Scale bars: 100µm. **d,** IHC confirming expression of HA-tag in NeuN^+^ neurons four weeks after initial labeling as DCX^+^ neuroblasts. NeuN^−^ cells in EPL and GL display mostly neuronal morphology. Most eYFP^+^-cells lost DCX expression indicating their progression to a more differentiated stage. GCL/MCL cells were mostly mature NeuN^+^-neurons (eYFP^+^/DCX^−^/NeuN^+^: 67.7%±2) or NeuN^−^-neurons (eYFP^+^/DCX^−^/NeuN^−^: 7%±1.7). Again, only few cells could be found in the EPL with similar characteristics compared to the three days timepoint (mostly eYFP^+^/DCX^−^/NeuN^−^: 3.1%±0.4). PGL cells were almost exclusively mature neurons (eYFP^+^/DCX^−^/NeuN^+^: 20%±2.5). n=3. Scale bars: 100µm. All quantifications are based on IHC depicted in Extended Data Figure 2. RMS: rostral migratory stream, dpi/wpi: days/weeks post induction, GCL: granular cell layer, MCL: mitral cell layer, EPL: external plexiform layer, PGL: periglomerular layer.

As opposed to previous studies that have used this system to study terminally differentiated cells^14,15,18^, targeting of cells along their maturation stages requires (i) sufficient incorporation of the tagged-RPL22 in functional ribosomes and (ii) no contamination of the target-maturation stage by other stages. Thus, we first confirmed that this strategy allows targeting of the maturation-stage of interest by immunohistochemistry for HA, eYFP and different cellular marker (Fig. 2b-d and Extended Data Fig. 2g-k). We further confirmed efficient incorporation of HA-tagged RPL22 into translating ribosomes and excluded potential changes in global protein synthesis levels in cultured NSCs from TiCRY mice (Extended Data Fig. 2a-f). Together, this characterization shows that careful selection of the reporter mouse line, timepoints and regions allows simultaneous ribosome-tagging and fluorescent-labeling of four distinct cell populations *in vivo* encompassing the complete adult OB-lineage progression: NSCs, ENBs, LNBs and neurons.

## RiboTag system allows parallel study of transcriptome and translatome

Next, to examine the transcriptome (total RNA) and translatome (Ribosome-bound RNA) we extracted the different RNA fractions from these maturation stages *in vivo*. Immunoprecipitation of HA-RPL22 allowed isolation of mRNAs bound to ribosomes (RIBO+). To correct for unspecific binding of mRNAs we included samples of mock immunoprecipitation of SVZ or OB tissue from Cre-negative animals (RIBO-). For isolation of total RNA of the same target population, we sorted eYFP^+^-cells (RNA; Extended Data Fig. 3b). First we validated enrichment of the expected population-specific marker genes in the RIBO+ and RNA fraction by quantitative RT-PCR (Extended Data Fig. 3a,c). Thereafter we profiled the RNA from the RNA, RIBO+ and RIBO-fractions by next generation sequencing. Sequencing results confirmed high enrichment for cell type specific transcripts both at the level of transcriptome (RNAseq) and translatome (RIBOseq)(Fig. 3 a,b; Supplementary Table 1). While NSC transcripts were mostly restricted to NSCs in the SVZ, neuroblast genes were already expressed at the level of NSCs (Fig. 3a). Similarly, expression of neuronal transcripts started already at the immature stage of LNBs (Fig. 3b). To further investigate the transcriptome and translatome, we performed differential expression analysis for stage transitions solely based on RNAseq or RIBOseq data, respectively (Fig. 3c-e; Extended Data Fig. 3d,e; Supplementary Tables 2, 3). Notably, the key core-regulators of neural development were represented in the transcriptome and translatome throughout the transition between the different differentiation stages (RNAseq and RIBOseq overlap) (Fig. 3c-e). Collectively, these data demonstrate that our system can be used to obtain high purity population (Extended Fig. 1c-e) and for integrated stage-specific transcriptome and translatome analysis.

**Figure 3.**
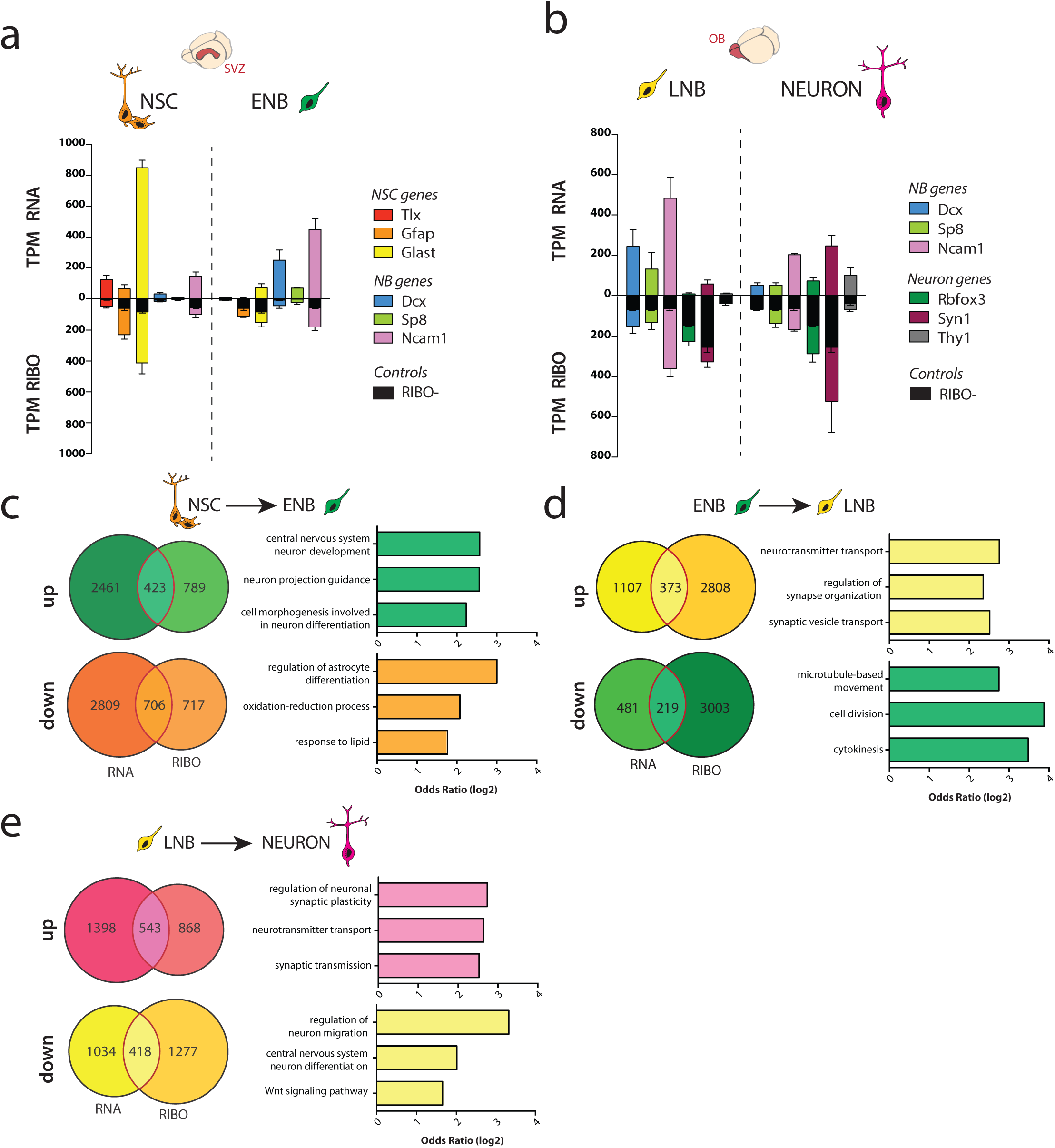
RiboTag system allows parallel study of transcriptome and translatome. **a,** Expression of stage-specific marker genes for SVZ populations at the level of transcriptome (top) and translatome (bottom). **b,** Expression of stage-specific marker genes for OB populations at the level of transcriptome (top) and translatome (bottom). Expression values based on tags per million (TPM) reads after Trimmed Mean of M-values (TMM) normalisation. Black bars in RIBOseq base on mock immunoprecipitation using Cre-mice (no HA-tag expression, RIBO-) representing average expression in the respective tissue corresponding to background binding (=noise). **c-e,** Overlap of differential expressed genes between RNAseq and RIBOseq for transition of NSC to ENB (c), ENB to LNB (d) and LNB to NEURON (e, FDR=10%). Only protein coding genes are considered. Gene set enrichment analysis on shared genes (marked in red) confirms enrichment of stage specific gene ontologies.

## Transcriptome and translatome diverge during lineage progression

Intriguingly, our analysis additionally revealed the existence of subsets of genes that are either less or more efficiently loaded onto ribosomes as compared to available mRNA (Fig. 3c-e). Besides, it showed a surprisingly low overlap between RIBO-seq and RNA-seq in the transition from ENB to LNB (Fig. 3d). This is the only comparison of cells extracted from two different regions, SVZ for NSC and OB for ENB, which highlighted the need of normalization. Thus, in order to identify transcripts, whose binding to ribosomes is post-transcriptionally controlled, we fit a linear model to explain the RIBOseq data by the RNAseq data (representing default translation) and the RIBO-controls (representing background binding) for each stage of interest (Fig. 4a). To identify targets with high confidence, we restricted our analysis to protein coding, highly abundant transcripts (read count > 1000 in RNAseq). Transcripts that were less abundant in RIBOseq compared to RNAseq (and RIBO-) were entitled „repressed“, suggesting that active mechanisms prevent them from ribosome association. In contrast, transcripts with higher ribosome association than expected based on RNAseq data and RIBO-data were entitled „enhanced“, indicating that their ribosome association is actively promoted. Our analysis revealed that abundant transcripts readily bind to ribosomes in NSCs (Fig. 4b,c,f and Supplementary Table 4), indicating little post-transcriptional regulation as previously reported in homeostatic HSCs^19^. Excitingly, with the onset of differentiation we observed a higher divergence of transcriptome and translatome (Fig. 4b,c,f, Extended Data Fig. 4a, and Supplementary Table 4). Neurons showed the highest degree of uncoupling between RNAseq and RIBOseq. Notably, most analyzed genes showed either repressed or enhanced translation as compared to their mRNA levels emphasizing the important role of posttranscriptional regulation for neuronal differentiation and function. The analysis of explained variance also confirmed that translation efficiency justifies a significantly higher fraction of the RIBOseq data in neurons as compared to RNA abundance (Extended Data Fig. 4b,c). We further examined the subset of transcripts sharing common translational control at the different maturation stages (Fig. 4d-f). First, we observed many transcripts exclusively repressed or enhanced at the ENB stage, some of which were validated via western-blot (WB) and qPCR (Extended Fig. 4 d-f). Second, LNBs and neurons exhibited the highest share of repressed transcripts, indicating that post-transcriptional repression of translation already starts at the LNB stage and is kept in neurons. Collectively, these data indicate that with differentiation to mature neurons, cells enter a mode of increasing dependency on posttranscriptional control.

**Figure 4.**
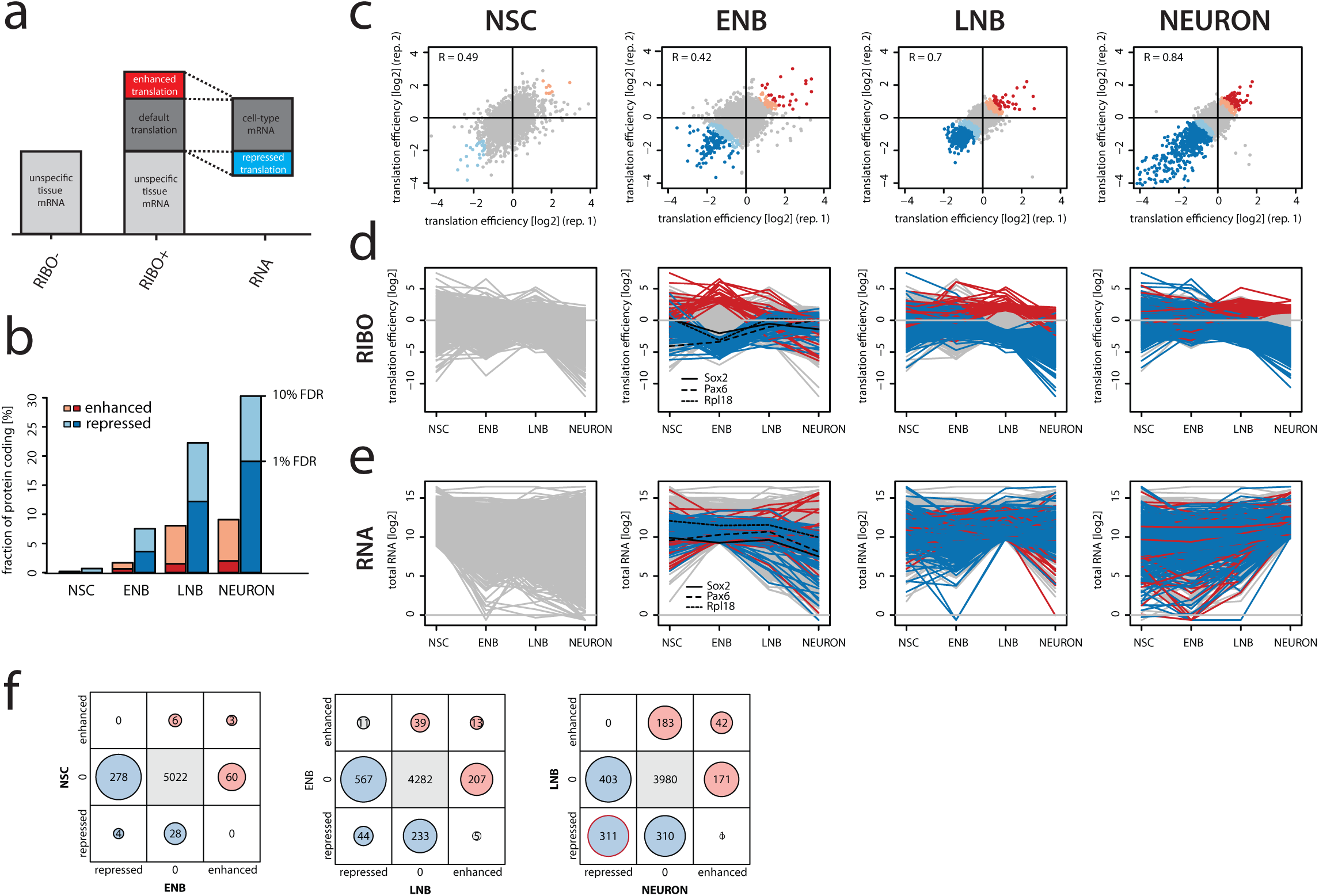
Stage-specific repression and enhancement of transcripts. **a,** Scheme describing the three datasets, which are implemented into analysis of translation efficiency. RIBO-sample resembles mock immunoprecipitation with HA-tag negative mice to assess background binding. **b,** Summary of the fraction of genes, which are repressed or enhanced in each population (only protein coding, >1000 reads). Few transcripts showed repression (32/4500, p=0.1) or enhancement (9/4500, p=0.1) in NSCs. ENBs feature a higher fraction of genes that were translationally repressed (282/3445, p=0.1) or enhanced (63/3445, p=0.1). LNBs displayed higher divergence of transcriptome and translatome (Repressed: n=622/1958, p=0.1; Enhanced: n=225/1958, p=0.1). Neurons showed the highest degree of variation between RNAseq and RIBOseq. Most of the analyzed genes showed repression (n=714/1443, p=0.1) or enhancement (n=214/1443, p=0.1). **c,** Scatter plots for each population showing translation efficiency. Grey dots: genes with linear ratio between transcript abundance (RNA) and translation (RIBO), blue dots: genes which are translationally repressed (dark: 1% FDR, light: 10% FDR), red dots: genes with enhanced translation (red: 1% FDR, orange: 10% FDR). Increasing number of genes with enhanced or repressed translation with further differentiation. **d,** Matched plots to the order of scatter plots in (b) illustrating temporal changes in translation efficiency of genes being significantly repressed or enhanced in the respective population. Only highly significant genes marked here in color (FDR=1%), Sox2 marked in black as an example for genes that are translationally repressed at the ENB stage. **e,** Matched plots to the order of scatter plots in (b) illustrating temporal changes in transcript abundance of genes being significantly repressed or enhanced in the respective population. Only highly significant genes marked here in color (FDR=1%), Sox2 marked in black as an example for genes that are translationally repressed at the ENB stage. **f,** Comparison of repressed- and enhanced genes over stages reveals increasing overlap with progression of cells. LNBs and neurons feature high overlap of repressed genes (n=311).

## Stage-specific repression of Sox2 translation

We next focused on the exit of the stem-state for transitioning into migrating ENBs. This is the first stage showing substantial control of ribosome binding of abundant mRNAs. Notably, the subset of repressed transcripts comprised multiple ribosomal genes as well as the prominent developmental transcription factors Pax6 and Sox2 (Fig. 4d). Sox2 and Pax6 have a highly evolutionary conserved role in determining the stem cell identity of embryonic and adult radial-glial progenitors together with Sox9^20^. In addition Sox2 is one of the factors used to reprogram somatic cells into pluripotent stem cells^21^. To validate Sox2 sequencing profile (Fig. 5a), we applied parallel *in situ* hybridization (ISH) for Sox2 mRNA and immunohistochemistry (ICC) for Sox2 and Dcx protein in the Tlx-reporter mice. This allowed simultaneous assessment of Sox2 mRNA and protein in NSCs and ENBs (Fig. 5b-d and extended Fig. 5c). While both NSCs and ENBs showed substantial levels of Sox2 mRNA, protein expression was mostly limited to NSCs (Fig. 5c,d). Interestingly, ENBs produce little Sox2 protein even when containing high levels of its mRNA. Comparison of protein expression to mRNA abundance revealed a 2.8-fold reduction of Sox2 translation efficiency from NSCs to ENBs. In addition, we independently confirmed that Sox2 shows highest translation efficiency in NSCs as compared to NPs and ENBs in freshly sorted SVZ cells via ISH and ICC but also via WB and qRT-PCR (Fig. 5e and Extended Data Fig. 5a-d). Interestingly, NPs, whose maturation state lies between NSCs and ENBs, show an intermediate efficiency for Sox2 translation (see regression lines), even if they displayed the highest absolute levels of Sox2 mRNA and protein. These data demonstrate a gradual repression of Sox2 translation in the transition from NSCs through NPs to ENBs. Interestingly, Sox2-mRNA regains binding to ribosomes later in LNBs and neurons (Fig. 5a). Thus, we further validated Sox2 translation in newborn neurons using DiCRY mice. While, adult born neurons in the granular cell layer did not show any Sox2 protein expression, periglomerular neurons exhibited high levels of Sox2 protein (Fig. 5f,g). Indeed, we identified Calretinin-positive newborn periglomerular neurons as the only neurons in the olfactory bulb expressing Sox2 protein (Fig. 5f,g). Thus, translation of Sox2 transcripts is transiently repressed in ENBs and becomes re-expressed in a subset of newborn olfactory bulb neurons. We next asked whether the set of 282 translationally repressed transcripts in ENBs would share distinct features as compared to non-repressed transcripts. *De novo* motif analysis on both untranslated regions (UTRs) as well as coding region of repressed transcripts discovered enriched motifs in the 5’ UTR including a stretch of pyrimidines (TCTTTC/CTCTTT, Supplementary Table 5) that we coined pyrimidine rich motifs (PRMs). A stretch of pyrimidines is also found in 5’ terminal oligopyrimidine (TOP) motifs, which are particularly sensitive to translation-mediated by mTORC1 activity^22^. Unlike the classical TOP motifs, this PRM is not always located at the terminal end of 5’UTR. Similarly to TOP-containing transcripts, PRM was found in transcripts encoding components of the translation machinery, such as ribosomal proteins (Fig. 6a, Extended Data Fig. 6a). Interestingly, PRM-containing transcripts were not repressed at the LNB stage despite the higher number of translationally repressed transcripts at this stage (Fig. 6b). Thus, we hypothesized that mTORC1 activity might be low in ENBs and high in NSCs and LNBs. This assumption is in conflict with the reported phosphorylation of 40S ribosomal protein S6 (rpS6), used as readout of mTORC1 activity, in proliferating neuroblast, but not in SVZ-neural stem cells^23^. In order to visualize phosphorylation of rpS6 specifically in NSCs and ENBs *in situ*, we used the Tlx-CreER^T2^-eYFP reporter mouse in combination with immunofluorescence staining for Dcx. Notably, phosphorylation of rpS6 (pS6) was low in all ENBs while NSCs showed heterogeneous levels of pS6, probably reflecting their different activation status (Fig. 6c). We further assessed expression of mTOR-related factors in freshly isolated ENBs and NSCs via WB. As predicted from the Ribo-seq analysis (see supplemental material), ENBs exhibited a downregulation of pS6 and phosphorylated-p70-S6K, and of total TSC2 and Rheb as compared to NSCs (Fig. 6d). In contrast, LNBs in the olfactory bulb showed significantly higher levels of pS6 than neigbouring neurons (Extended Data Fig. 6f). Interestingly, neurons of the mitral cell layer in the olfactory bulb showed high pS6 levels (data not shown), suggesting variability between different neuronal subtypes regarding mTOR activity, which might translate into protein synthesis rates. Thus, the drop of mTORC1 in ENBs but not LNBs could be the reason for the specific repression of PRM-transcripts in ENBs. To experimentally validate the role of mTORC1 in translation of the PRM-transcript Sox2 *in vivo*, freshly isolated ENBs were acutely treated with membrane permeant phosphatidylinositol 3,4,5-trisphosphate [PI(3,4,5)P3]. This lipid is produced upon nutrients and growth factor activation of PI3K and mediates AKT activation of mTORC1^24^. ENBs were either treated with PI(3,4,5)P3 alone or together with the mTOR inhibitor Torin1, which unlike rapamycin fully inhibits mTORC1^22,25^. Thereafter, levels of pS6 and Sox2 proteins were assesed by ICC in single cells. First, this assay confirmed that NSCs exhibit higher levels of pS6 and Sox2 than ENBs (Fig. 6e). Second, exposure of ENBs to PI(3,4,5)P3 highly increased pS6 levels in ENBs to levels comparable to NSCs (Fig. 6e). PI(3,4,5)P3-mediated pS6 increased could be inhibited by Torin. Most importantly, PI(3,4,5)P3 also increased Sox2-protein levels in a mTORC1-dependent manner, as Sox2 protein expression could be repressed by Torin (Fig. 6e). Next, we wanted to examine the mTORC1-dependent rate of ribosomal load of PRM+ and PRM-transcripts. To this end we turned to NSCs cultures to increase the input material and be able to gain cell-fractions with distinct ribosome content via sucrose gradient fractionation of Torin1- or vehicle-treated NSCs. Most PRM+ transcripts but not PRM-ones, shifted to the lighter fractions upon Torin1-treatment, indicating that their translation depends on mTORC1 activity (Extended Data Fig. 6d). Surprisingly, this shift could not be detected for Sox2-transcripts (Extended Data Fig. 6d,e). Therefore we compared the expression of mTORC-related factors in cultured NSCs to expression in freshly isolated NSCs and ENBs. In cultured NSCs were left untreated or treated with Torin or the EGFR inhibitor, erlotinib. The latter was used to mimick the loss of EGFR occurring in NSCs transitioning to ENBs. Both Torin and Erlotinib treatment of NSCs *in vitro* led to a transient reduction in pS6 and phosphorylated-p70S6K, but not to lowering expression of the mTORC1-activator Rheb as found between freshly isolated ENBs and NSCs (Fig. 6d and Extended Fig. 6 b,c). These findings suggest that lineage transitions induced stoichiometric changes that cannot be achieved by an acute inhibition of mTORC1. Surprisingly, Torin-treated cultured NSCs exhibited a slight upregulation of protein expression of genes exhibiting enhanced translation efficiency in the transition from NSCs to ENBs *in vivo* (Extended Fig. 6 e). These genes are overrepresented in the neuron-differentiation-related gene ontology categories (Extended Fig. 6a). Thus, these data suggest that a drop of mTORC1 orchestrates posttranscriptional repression of stemness and enhancement of neuronal differentiation processes. Along this line, long-term activation of mTORC1 via deletion of TSC2 in human embryonic stem cells generated neurons with increased expression of neural stem cell and proliferation markers including SOX2, Nestin and PDNA and decreased expression of neuronal markers^26^. Altogether these data underline the importance of studying the molecular underpinnings of lineage transitions *in vivo* and unveils a yet unrecognized differential sensitivity of targets to long-term and short-term mTORC1 inhibition that needs to be addressed in future studies.

**Figure 5.**
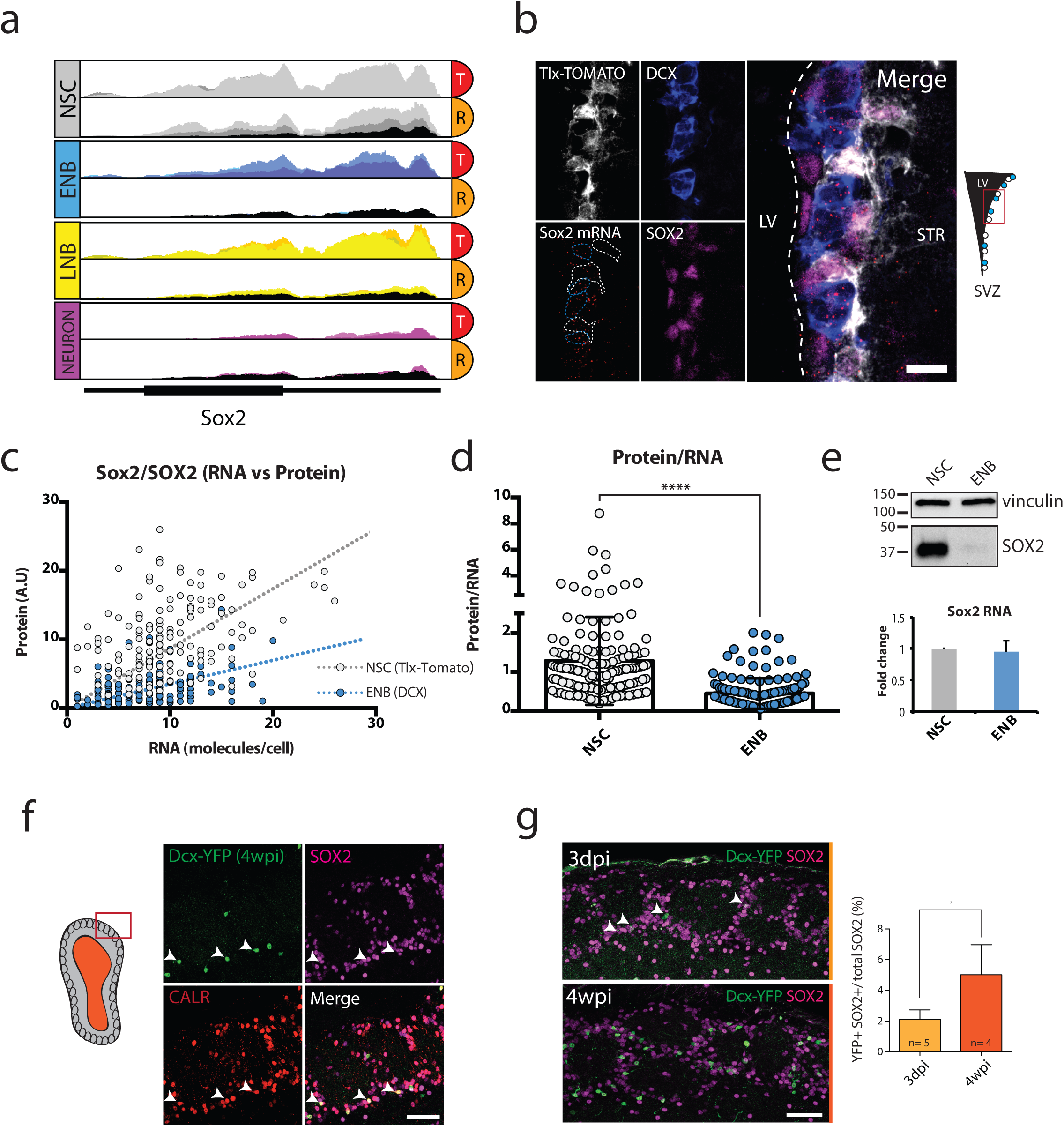
Sox2 is post-transcriptionally repressed in ENBs and expressed at high levels in a subpopulation of olfactory bulb interneurons. **a,** Sox2 locus in the integrated genome viewer (IGV) both of total RNA (T) and ribosome-bound RNA (R) for populations of interest, histogram shows abundance of reads. Different color hues mark biological replicates. Black histogram indicates background binding (RIBO-). Sox2 shows no ribosome binding specifically at ENB stage indicating post-transcriptional repression. **b,** Representative image for parallel *in situ* hybridization and immunohistochemistry for Sox2 protein/RNA in NSCs (marked by Tlx-TOMATO) and ENBs (marked by DCX) validating Sox2 repression event. Scale bar: 10µm. **c,** Quantification or Sox2 RNA (molecules per cell) and protein expression (total fluorescence) by manual segmentation. Regression line indicates different translation efficiency. Quantified 144 NSCs and 145 ENBs from two mice. **d,** Ratio of protein expression to RNA abundance indicates greatly reduced translation efficiency of SOX2 in ENBs. Significance by students t-test (Mann-Whitney). **e**, Sox2 protein and mRNA abundance in freshly sorted NSCs and ENBs. Protein levels were analyzed by WB (representative WB, upper panel) and the mRNA was quantified by qPCR (lower panel). Vinculin served as a loading control. The fold change was calculated based on three technical replicates. **f,** Staining of olfactory bulb periglomerular layer showing Dcx-eYFP traced cells (4wpi) which coexpress SOX2 and the neuronal marker Calretinin. Scale bar: 50µm. **g,** Comparison of short (3dpi) and long (4wpi) tracing of Dcx-eYFP cells shows increase of eYFP^+^SOX2^+^ cells with time indicating contribution of migrating, newborn neurons to the pool of SOX2^+^ neurons in the periglomerular layer. Scale bar: 50µm.

**Figure 6.**
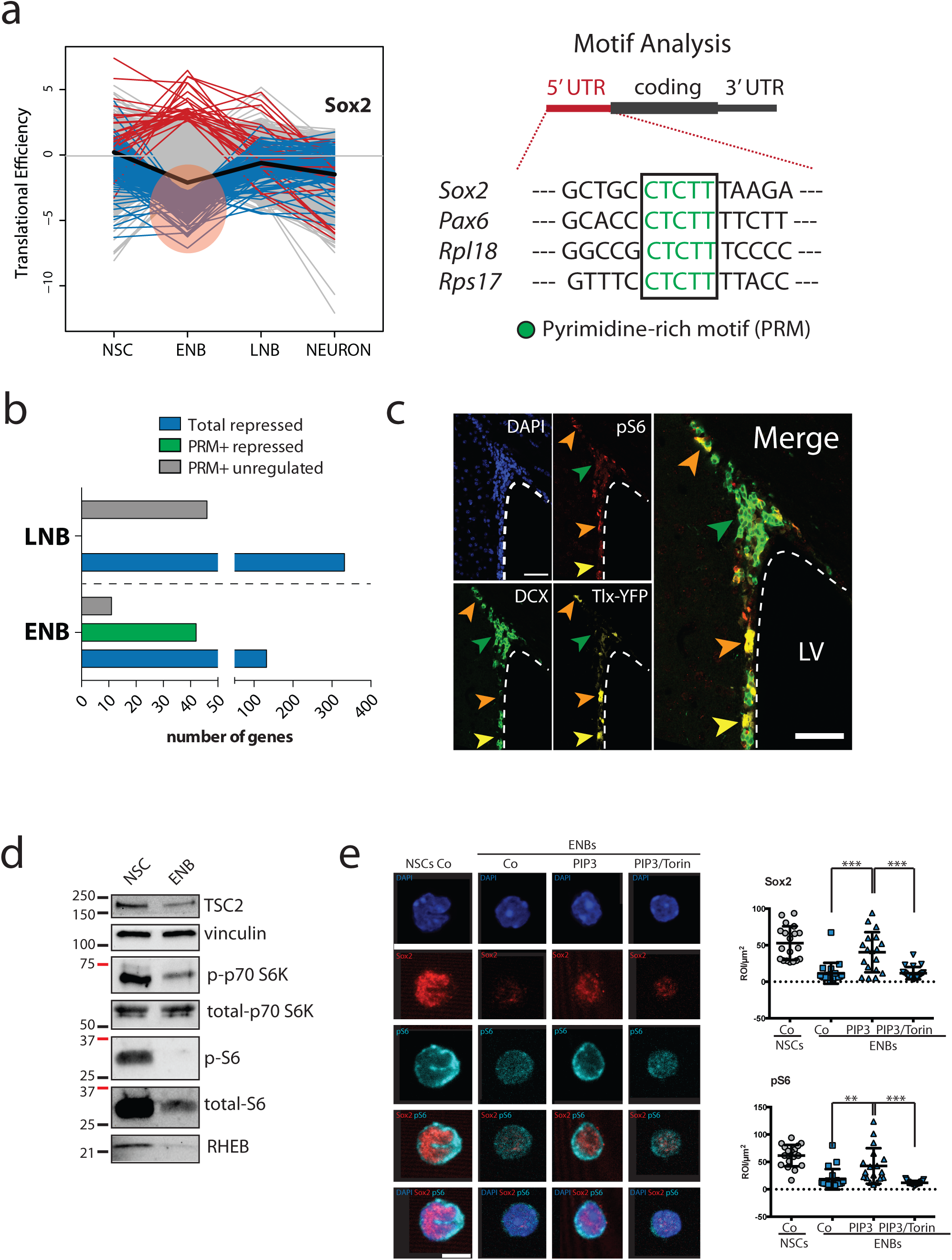
Pyrimidine-rich motifs (PRMs) in the 5’ UTR predict translational repression under mTORC1 mediated global arrest. **a,** Motif analysis reveals enrichment of pyrimidine-rich motif (PRM) CTCTT in the 5’ untranslated region (UTR) of genes, which are repressed at the ENB stage. **b,** Relation of the presence of a 5’ PRM and a drop in translation efficiency at the ENB or LNB stage indicates stage-specific repression of 5’ PRM containing transcripts in ENBs independent of the total number of repressed genes (1% FDR). **c,** Staining for phosphorylation of ribosomal protein S6 (pS6) in the SVZ as readout for mTORC1 activity. Comparison of NSCs (Tlx-eYFP^+^) and ENBs (DCX^+^). All ENBs show low levels of pS6 (green arrow), while some NSCs show high pS6 (orange arrows) and others show low levels of pS6 (yellow arrow). Scale bar: 50µm. d, WB analysis of the steady-state level of a number of proteins involved in mTOR signaling in NSCs and ENBs freshly sorted from the SVZ. Vinculin served as a loading control. **e,** Freshly isolated ENBs were directly fix or exposed to PIP3 alone or with Torin and the levels of pS6 and Sox2 assessed by ICC. NSCs were directly fix after isolation and stained by ICC. Representative images of Sox2 (red) and pS6 (turquoise) in the different groups are shown (left panel). Scale bar: 5µm. The relative expression of Sox2 and pS6 in the different groups is shown in right panel. Statistical significance was calculated by students t-test p < 0.01^**^, p < 0.005^***^.

Thus, our data uncovers a way of regulation of components of the translation machinery and neurogenic transcription factors via modulation of mTORC1-activity in the natural environment of the brain. An activation of mTORC1-activity is found in injured muscle stem cells, which transit from a quiescent (G0) to a quiescent primed state to finally enter cell cycle (G alert)^11^. Yet the specific transcripts underlying mTORC1-control of G alert in muscle cells remain unknown. Most importantly, this might represent a more general mechanism by which mTOR controls stemness.

## DISCUSSION

Altogether, this study demonstrates that the rate of active protein synthesis is tightly controlled during stem cell differentiation into neurons. This general activation of protein synthesis is required to exit the dormant state in NSCs^2^ and muscle stem cells^11^ and to activate differentiation in skin stem cells^8^ and drosophila germline stem cells^27^. Notably, impaired drop of mTORC1 by deletion of TSC1 in single SVZ-NSC in the rodent brain, led to generation of Dcx-negative cells with immature radial-like morphology that were unable to migrate out of the SVZ^28^. Our data indicates that the observed impaired neurogenesis is due to loss of translation arrest in ENBs needed to repress stem cell identity prior to a re-establishment of protein synthesis in LNBs to allow neuronal maturation. There has been long-standing discussions about the overall correlation of transcript- and protein abundance with contradicting conclusions from identical datasets^29,30^. However, with the forthcoming of refined mathematical models and latest technologies for quantification, it is believed that mRNA levels largely explain protein abundance during steady state^31^. Notably, the study of developmental lineage transitions *in vivo*, has allowed us to uncover a highly dynamic and stage-specific regulation of translation efficiency. Whereas the transcriptome and translatome highly correlate in stem cells during homeostasis, they become increasingly divergent with the exit of the stemness state. The relatively large number of repressed mRNAs in our system is not surprising if we remember that stem cells partially recapitulate some features of germ cells, and the latter rely considerably on translational regulation instead of transcriptional during most of their lifespan. Furthermore, recent data revealed embryonic origin of adult NSCs^6,7^. The stock of mRNAs in the cytoplasm, stable but not translated, allows the cell to respond quickly to stimuli. NSCs rearrange their morphology, contacts with the niche and migrate to a new niche upon their differentiation. Thus, for such cells relying on post-translational regulation might be crucial to be able to quickly adapt to fast-changing environment conditions. Importantly, selective changes in translation efficiency have been reported in oncogenic NSC in a spontaneous brain tumor model^32^.

In early neuroblasts, a drop in mTORC1 activity programs a transient arrest in translation of PRM+ transcripts, which include ribosomal genes and key evolutionary conserved determinants of stem cell identity. Exit of stemness requires tight regulation to ensure continuos production of differentiated progeny while maintaining the pool of stem cells. Dysregulation of this transition is found in aging and cancer^33^. mTOR is hyperactivated in 90% of glioblastomas^34^. Notably, mTOR inhibition by rapamycin in glioblastoma abrogated expression of Sox2 protein thereby reversing their chemoresistance to temozolomide^35^. Thus, this study uncovers an unrecognized role of mTOR in stem cells that together with activation of a G-alert state and proliferation contributes to explain the dependency of cancer stem cells on mTOR activity. Besides, we identify a biological process in the adult brain *in vivo* that critically depends on the selective sensitivity of PRM+ transcripts to mTOR-mediated translation. Likewise, despite loss of mTORC activity, a subset of mRNAs increased their translation efficiency. The main effect of mTOR on translation is the phosphorylation and inactivation of 4EBP proteins, which associate and inactivate a cap-binding initiation factor eIF4E responsible for 40S subunit recruitment to mRNAs. Those mRNAs, showing high dependence on the cap structure and eIF4E, are among the most sensitive to mTOR activity. Many such mTOR-sensitive mRNAs encode the most abundant proteins in cells, including ribosomal proteins. These mRNAs are very efficient in translation and form a class of so-called “strong” mRNAs^36^. However, the advent of the next-generation sequencing technology revealed a considerable number of mRNAs, which show resistance to mTOR repression and even get stimulated^22,37^. Additional studies found out that many such mRNAs are less dependent on eIF4E and can apply other mechanisms of 40S recruitment independent on the eIF4E-cap interaction^38^. These mRNAs could use isoforms of initiation factors eIF4E and eIF4G, which do not bind 4EBP repressors and could confer less dependence on mTOR to the corresponding mRNAs. Furthermore, these mRNAs being less competitive upon optimal conditions get considerable advantages upon the repression of “strong” mRNAs, relishing uncompetitive availability of ribosomes, tRNAs, and translation factors. Although, the specific elements of such mRNAs providing mTOR independence are not yet well described, recently discovered capability of N6-methylated adenines localized in the 5’-UTRs to directly attach 40S in an eIF4E-independent manner might be one of those^39^. Our motif analysis solely revealed enrichment in the coding region of a consensus motif for m6A “CGCAAC” (as shown in table S5), though this is not sufficient to explain escape from mTORC1 inhibition. Future studies of N-6-methylated adenines should further our understanding of increased translation efficiency in the frame of low mTORC1 activity for this set of transcripts.

In summary, our data offers a comprehensive genomic view on translational control during neuron generation in the adult brain, adding a novel layer to our knowledge of the molecular mechanisms controlling stem cell differentiation.

## Acknowledgements

We thank S. Wolf from the DKFZ Genomics and Proteomics Core Facility; V. Eckstein from the Heidelberg University Hospital FACS Core Facility; Monika Langlotz from the ZMBH FACS Core Facility, Hai-Kun Liu for Tlx-CreER and tdTomato-flox mice; Markus Schwaninger for Dcx-CreER^T2^ mice; David Wiest for the Rpl22 antibody; Klara Zwadlo, Stefanie Limpert and Katrin Volk for technical assistance; Damir Krunic and the Light Microscopy Core Facility for support with image analysis; Bernd Bukau and Günther Kramer for advice during the early stage of the project, the TAC members Aurelio Teleman, Georg Stöcklin and Matthias Hentze and the members of the Martin-Villalba laboratory for critical comments. We specially thank Aurelio Teleman for critical reading of the manuscript. This work was supported by the University of Heidelberg and DKFZ (bridge-project ZMBH-DKFZ alliance), the DFG (SFB873), the BMBF and the DKFZ.

## Author contributions

Project concept, A.M.-V, A.B.; Investigation, AB, Y.D., M.S., G.S.G.B., G.K., S.K., M.G., A.S.L., E. LL.B., R.S., B.F.; Writing-Review & Editing, A.M.V., B.F., A.B.; Supervision, A.M.-V., and B.F.; Project Oversight, A.M.V., B.F.

## Author information

RNAseq, RIBOseq and RIBO-raw sequence data is available from GEO (Accession number: GSE94991). Complete documented software is available as supplementary file. The complete R/Bioconductor software package will be made available upon acceptance on the authors’ webpage (https://martin-villalba-lab.github.io/). The authors declare no competing financial interests. Correspondence and requests for materials should be addressed to AMV.

## METHODS

### Mice

Male transgenic mice of DiCRY (Dcx-CreER-Rpl22.HA-eYFP) and TiCRY (Tlx-CreER-Rpl22.HA-eYFP) lines were used between 8 and 12 weeks of age for all sequencing experiments. These mice were generated by mating of RiboTag mice (B6N.129-Rpl22tm1.1Psam/J, purchased from The Jackson Laboratory) with Tlx-CreER-eYFP mice^40^ or Dcx-CreER-eYFP mice^41^. Additionally, TiCROMATO mice (Tlx-CreER-Rpl22.HA-Tomato) were used whenever eYFP expression was considered to be too weak. All mice were backcrossed to a C57BL/6 background.

All further experiments using transgenics were conducted using both male and female mice up to 16 weeks of age. Recombination was induced by oral gavage of tamoxifen (TAM, Sigma-Aldrich) at a concentration of 200mg/kg body weight (volume ~100µL; solved in 90% sunflower oil/10% EtOH). TAM was administered twice per day for a total of 2-3 injections and the experiments were conducted 3-4 days after recombination, since ribosome turnover in the brain takes several days^42^. The conditions varied slightly between mouse lines due to recombination efficiencies and total number of cells.

C57BL/6 male mice between 8 and 12 weeks of age were used for all other experiments. All mice were bred inhouse at the DKFZ Center for Preclinical Research. Animals were housed under standard conditions and fed ad libitum. All procedures were in accordance with the DKFZ guidelines and approved by the Regierungspräsidium Karlsruhe.

### Cell sorting

Animals were sacrificed by cervical dislocation and brains were immediately placed on ice. The lateral SVZ was microdissected as a wholemount as previously described^43^, further the olfactory bulb were dissected. Tissue of 1-2 mice was used per replicate (SVZ or OB) for sequencing experiments. Tissue of up to five mice was pooled per sample for OPP experiments. Tissue was digested with trypsin and DNase according to the Neural Tissue Dissociation kit in a Gentle MACS Dissociator (Miltenyi). For sequencing experiments endogenous eYFP fluorescence was used to identify cells of interest. For OPP experiments cells were sorted as previously described^2^ using following antibodies: GLAST (ACSA-1)-PE (Miltenyi, 1:20), APC-Cy™ 7 CD45 (BD, 1:200), O4-APC (Miltenyi, 1:50), Prominin PerCP-eFluor710 (eBioscience, 1:75), PSA-NCAM-PE-Vio770 (Miltenyi, 1:50), conjugated EGF-Alexa488 (Life Technologies, 1:100). During cell sorting, Sytox blue (Thermo Fisher Scientific, 1:1000) was used to exclude dead cells. For transcriptome analysis, 200-500 cells were sorted directly into Picopure lysis buffer (Thermo Fisher Scientific). Sequencing libraries were prepared as described in the following section for immunoprecipitation samples.

### RiboTag immunoprecipitation and sequencing

RiboTag immunoprecipitation was conducted essentially following the original protocol^15^ with minor modifications. Animals were perfused with Hank’s Balanced Salt Solution (HBSS) supplemented with the translation inhibitor cycloheximide (CHX, 200 µg/ml) for stabilization of RNA-ribosome complexes. Tissue of interest was dissected and placed in tubes containing 1.4mm ceramic beads (PEQLAB). Homogenization buffer (50mM Tris HCl, 100mM KCl, 12mM Mg_2_Cl, 1% NP-40, 1mM DTT, 1x Roche complete protease inhibitor, 200U/mL RNAsin from Promega, 100µg/mL CHX, 1mg/mL Heparin) was added to a weight per volume ratio of 3% (e.g. 0.015g in 500µL) and tissue was homogenized using Minilys Personal Homogenizer (Bertin Instruments). Samples were centrifuged and supernatant was used for immunoprecipitation. Anti-HA antibody (Covance, 1:100) was added for four hours (indirect conjugation) following addition of Protein G magnetic beads (100µL, prewashed with homogenization buffer, Thermo Fisher Scientific) and overnight incubation (all steps at 4°C). Supernatant was discarded and beads were washed three times for ten minutes in high salt buffer (50mM Tris HCl, 300mM KCl, 12mM Mg_2_Cl, 1% NP-40, 1mM DTT, 100µg/mL CHX). Finally, beads were resuspended in Picopure lysis buffer (supplemented with β-mercaptoethanol, 10µL/1mL, Sigma) and RNA was isolated by Picopure RNA isolation kit (Thermo Fisher Scientific) following the manufacturer’s protocol. For enrichment analysis, fraction of the RNA was converted to cDNA using the SuperScript VILO kit (Thermo Fisher Scientific). Sequencing libraries were prepared using Smart-seq2 technology as previously described^44^. Reverse transcription was performed using an oligo (dT) primer and a locked nucleic acid (LNA)-containing template-switching oligonucleotide (Exiqon). Full-length cDNAs were amplified by 15-18 cycles of PCR using KAPA HiFi DNA polymerase (KAPA biosystems). cDNAs were then converted into libraries for Illumina sequencing according to the Nextera XT Sample Preparation (Illumina) protocol. Samples were sequenced in an Illumina HiSeq 2000.

### Cell culture

For NSC isolation, 8-12 week old wild type C57BL/6 or TiCRY mice (one week post induction) were sacrificed by cervical dislocation. A wholemount of the SVZ was microdissected^43^ and cells were isolated using trypsin (or papain) and DNAse according to the Neural Tissue Dissociation kit in a Gentle MACS Dissociator (Miltenyi). NSCs were cultured in Neurobasal medium (Thermo Fisher Scientific) supplemented with 20 ng/ml of basic fibroblast growth factor bFGF and epidermal growth factor EGF.

Freshly sorted cells for OPP administration (translation analysis) or RNA *in situ* hybridization were plated in Poly-D-lysine- and laminin-coated Lab-Tek chambers (Thermo) in Neurobasal medium lacking growth factors. OPP was added to the culture (50µM) two hours after plating and one hour after incubation cells were fixed for 20min using 2% paraformaldehyde (PFA) in the medium. For RNA *in situ* hybridization cells were fixed for 20min using 2% PFA in the medium two hours after plating.

### Erlotinib, Torin1 and PI(3,4,5)P3 treatment

Primary NSCs were exposed to erlotinib (5 µM, stock in DMSO; SIGMA-Aldrich) or Torin1 (250 nM, stock in DMSO) for the indicated time periods. The solvent DMSO was used as a vehicle control. Thereafter cells were collected in lysis buffer and analyzed by western blot.

For the PI(3,4,5)P3 (PIP3) treatment, NSCs and ENBs were sorted from SVZ and either resuspended in Neurobasal media with the growth factors or fixed with 2% PFA. Thereafter, ENBs were either treated with PIP3 or PIP3/Torin1. Torin1 (250nM) treatment was done for 5 min prior to PIP3 stimulation. PIP3/AM(DOG) synthesized as previously described^45^, was prepared as a stock solution of 50mM in DMSO and stored at −80°C. Just before used PIP3 was mixed with 20% pluronic F-127/DMSO solution (Invitrogen) in a 1:0.5 ratio to facilitate cell entry. The PIP3/AM/pluronic/DMSO mixture was resuspended in 50µl ENB-medium to a final concentration of 10µM and immediately added to cells for 10minutes. Thereafter, PIP3 was removed and fresh medium and additional Torin1 was added for further 2hrs. The control sample was treated with DMSO as a vehicle control. Cells were fix and further process for immunocytochemistry (ICC).

### Western Blot analysis and qPCR

Around 10^4^ freshly sorted NSCs and ENB cells were collected in PBS with 10% FBS on ice for protein analysis by western blot. For re-probing, membrane stripping was performed according to the protocol from Cell Signaling technologies. For primary cultures, cells were collected, washed with PBS and lysed as described before.

The following primary antibodies were used for western blot analyses. Phos EGFR Tyr1068 (rabbit, 1:1000), EGFR (rabbit, 1:1000), phos S6 Ser240/244 (rabbit, 1:1000), S6 (rabbit, 1:10,000), phos p70 S6K Thr421/Ser424 (rabbit, 1:1000), p70 S6K (rabbit, 1:1000) and Tuberin/TSC2 (rabbit, 1:1000) were from Cell Signaling Technologies; DUSP4 (rabbit, 1:800), RHEB (mouse, 1:1000), RPS20 (rabbit, 1:1000), Sox2 (rabbit, 1:1000), VASH2 (rabbit, 1:1000), vinculin (rabbit, 1:5000) were from Abcam; actin (rabbit, 1:1000) and RPL22 (mouse, 1:1000) were from Santa Cruz; Sp8 (rabbit, 1:1000) was from Millipor; PLP1 (chicken, 1:400) was from Neuromics; anti-HA (mouse, 1:1000) was from BioLegend.

For qPCR analysis of mRNA abundance in freshly sorted NSCs and ENBs, we collected around 5,000 cells of each type. For primary NSCs in culture, around 10^5^ cells were used. Total RNA was extracted with the Picopure RNA isolation kit (Thermo Fisher Scientific). cDNA was synthesized using the SuperScript VILO kit (Thermo Fisher Scientific). qPCR reactions were performed in three technical replicates with the Power SYBR™ Green PCR Master Mix (Thermo Fisher Scientific) and QuantiTech primers specific for mouse genes (Qiagen). Cycles of amplification were made in the CFX384 Real-Time System (Bio-Rad). Excel (Microsoft) was used to analyse the qPCR data.

### Immunocytochemistry, immunohistochemistry and RNA *in situ* hybridization

Cells were fixed for 20min at room temperature using 2% PFA in the medium. Staining was performed using standard procedures with following antibodies: anti-Sox2 (mouse, Abcam, 1:500) and anti-pS6 Ser240/244 (rabbit, Cell Signaling, 1:1000). OPP visualization was done after all staining steps. For this, the alkyne group of OPP was detected using an Alexa488- or Alexa647-coupled azide according to the Click-iT Cell Reaction Buffer Kit (Life Technologies).

For tissue staining, mice were transcardially perfused first with HBSS and then with 4% PFA, following overnight postfixation. Vibratome sections of 50-70µm were prepared for tissue staining only, for RNA *in situ* hybridization cryosections of 11-13µm were prepared. Following antibodies were used for ICC/IHC: anti-DCX (goat, Santa Cruz, 1:500), anti-DCX (guinea-pig, Merck Millipore, 1:1000), anti-pS6 Ser240/244 (rabbit, Cell Signaling, 1:1000), anti-NeuN (mouse, Merck Millipore, 1:500), anti-pH3 (mouse, Merck Millipore, 1:500), anti-GFAP (mouse, Merck Millipore, 1:1000), anti-GFAP (rabbit, Merck Millipore, 1:1000), anti-HA (mouse, Covance, 1:1000), anti-GFP (chicken, Aves, 1:1000), anti-RFP (rabbit, Rockland, 1:1000), anti-Sox2 (goat, Santa Cruz, 1:500), anti-Sox2 (rabbit, Abcam, 1:1000),anti-Calretinin (rabbit, Merck Millipore, 1:1000), anti-GLAST (guinea pig, Frontier, 1:1000), anti-Mash1 (rat, RDI Fitzgerald, 1:500).

RNA *in situ* hybridization was performed using RNASCOPE Multiplex Fluorescent Assay (Advanced Cell Diagnostics) with one major modification of the protocol: To allow subsequent protein staining we skipped pretreatment steps as it is described in the manufacturer’s protocol and instead only did extended (15min, 95°C) heat-based target retrieval using a modified citrate buffer (Target Retrieval Solution, Dako). Additional immunofluourescence was done following the mRNA detection following standard procedures.

### Polysome fractionation and protein/RNA isolation

Polysome fractionation for cultured cells with subsequent protein and/or RNA isolation was performed essentially following a previously published protocol for CNS tissue leaving out initial steps of tissue preparation^46^. Cells were acutely treated for 5 min with cycloheximide in culture medium (100µg/mL) and subsequently collected, washed in PBS/CHX and lysed in polysome lysis buffer (PLB, 20 mM Tris HCl, pH 7.4, 5 mM MgCl_2_, 120 mM KCl, 1% Nonidet NP40, 100µg/mL CHX, 1x Roche complete protease inhibitor, 200 U/mL RNAsin from Promega, 14 mM b-mercaptoethanol). Lysates were centrifuged and the supernatant was added on sucrose gradients (17.5-50%). Samples were centrifuged for 2.5 hours at 35.000rpm at 4°C in an ultracentrifuge (Beckman L8M). Samples were fractionated in 12 fractions containing one mL sample using an Isco fractionator while monitoring absorption at 254nm wavelength. RNA isolation was performed using acidic Phenol-Chloroform-Isoamylalcohol (PCI, 25:24:1, Thermo Fisher Scientific). Alternatively, proteins were precipitated using trichloroacetic acid (TCA).

### Imaging

Confocal images were acquired using a Leica TCS SP5 confocal microscope. Image J was used for processing and images were only adjusted for brightness and contrast. For OPP quantifications, all images within compared groups were acquired with identical settings. Analysis was performed in ImageJ software using a custom-written macro for unbiased segmentation and quantification of pixel intensity and cell size. OPP quantifications are depicted as the integrated pixel intensity within a cell and normalized to a reference group within the same experiment. In graphs, mean and SD are shown. Illustrator CS5 (Adobe) was used for figure assembly.

### Processing and analysis of RNAseq and RIBOseq data

Sequence reads from both RNAseq and RIBOseq samples were aligned to the mouse reference genome (ENSEMBL Release 80) and the ENSEMBL gene annotation (Release 80) using the STAR alignment algorithm version 020201^47^ with the proposed the ENCODE settings to generate gene-specific raw count values. Expected gene counts and TPM values (transcripts per million) were computed using RSEM (version 1.2.21)^48^ with bowtie2 (version 2.2.6) with the same genome and verison and annotation. Differential gene expression analysis between developmental stages of the RNAseq data was conducted by the R/Bioconductor package DESeq2^49^. Thereby p-values were computed by fitting a negative binomial distribution and subsequent testing by a Wald test. P-values were corrected for multiple testing by controlling the false discovery rate applying the method of Benjamini-Hochberg. For RIBOseq data the same analysis was applied to analyze the differential abundance of ribosome bound RNA.

To assess the translation efficiency, genes with an average log2 read count of at least 10 were considered. At first, a linear model was fitted for each replicate individually to explain the RIBOseq data by a linear combination of RNAseq data (representing efficient translation) and RIBO- (representing tissue background expression). Analysis of variance of this part provides the proportion of variance that is explained by RNAseq and by RIBO- in each replicate separately. The RIBOseq signal that cannot be explained by RNAseq or RIBO- can be explained by either enhanced or repressed translation. To assess, if the detected enhanced or repressed translation was statistically significant, we compared the two replicates and test, if the remaining signal is different from zero by applying a moderated t-test implemented in the R/Bioconductor package limma^50^. P-values were corrected for multiple testing by the method of Benjamini-Hochberg. At this stage the analysis of variance provides the proportion of variance that can be explained by translation efficiency (correlated signal) and the remaining unexplained variance (uncorrelated signal) that is most likely of technical source. Full documentation of computational analysis can be found as a supplementary document.

### *De novo* motif analysis

3’ UTR, 5’UTR and protein coding sequences of all protein coding genes were tiled in 6-mer nucleotide sequences. For each developmental stage separately, for translationally repressed and enhanced genes separately, and for 3’UTR, 5’UTR and coding sequence separately, an unbiased motif analysis was performed. For each possible 6-mer, the number of genes containing the tested 6-mer in the set of repressed (enhanced) genes and in the remaining genes was computed. A p-value was computed by a binomial test and corrected by the method of Benjamini-Hochberg. All over- or underrepresented motifs at a false discovery rate of 10% were reported (Supplementary Table 5).

---

**Extended Data Figure 1.**
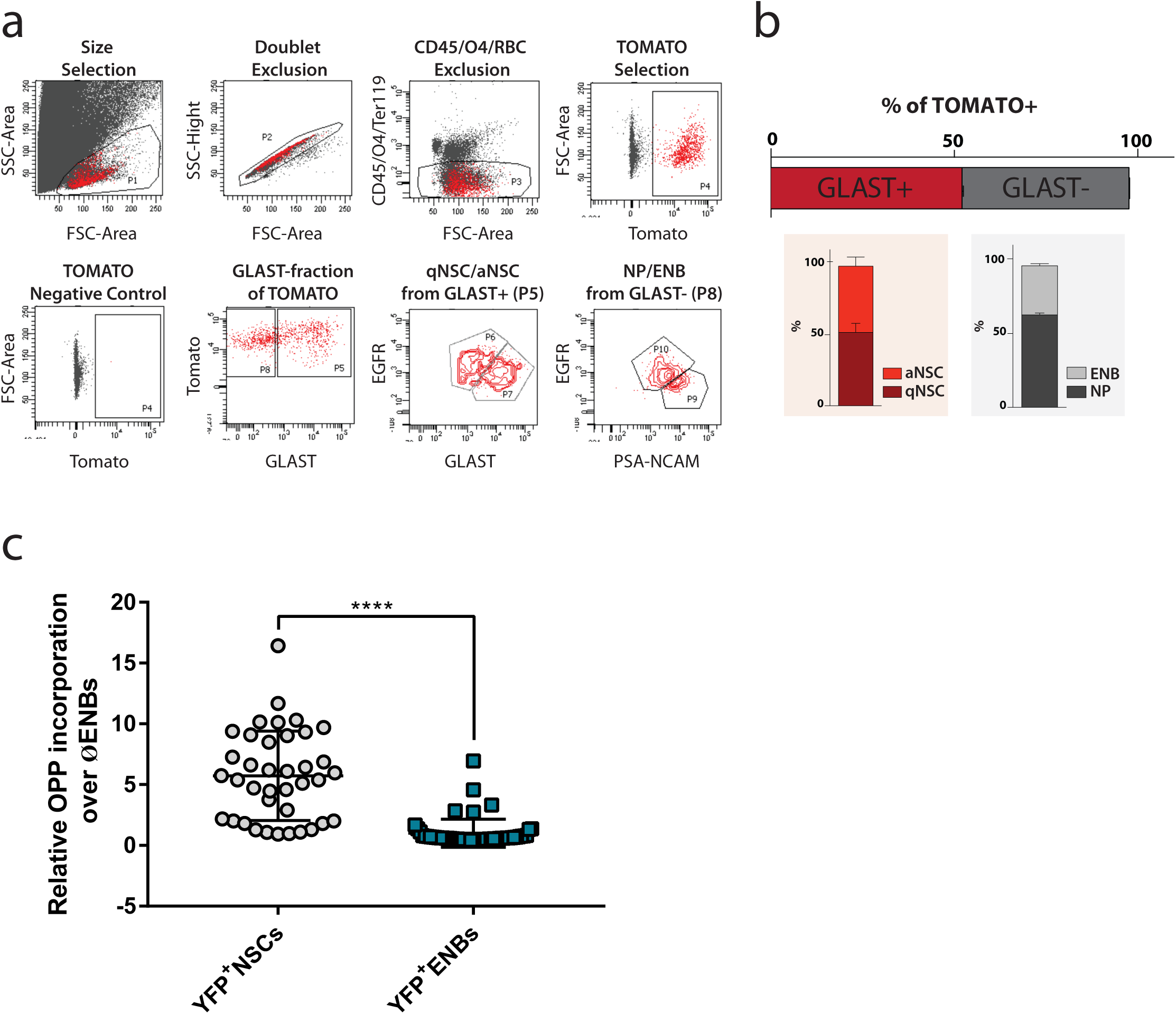
**a,** FACS gates for NSCs labeled by Tlx-TOMATO four days post induction. NSCs, which are labeled by recombination under the Tlx-promoter, contain both quiescent (GLAST^+^EGFR^−^) and active NSCs (GLAST^+^EGFR^+^). Prom1 was not used in this experiment since it is known that Tlx does not recombine in astrocytes. **b,** Composition of TOMATO^+^ cells four days after induction. 52.1%±0.6 are NSCs (GLAST^+^) while 45.6%±0.3 are non-NSCs (GLAST^−^). The numbers do not sum up to 100% since some few cells lay between gates. NSCs contain 51.7%±9.1 quiescent NSCs (EGFR^−^) and 46.7%±9.1 active NSCs (EGFR^+^). GLAST^−^ cells contain 62.7%±1.6 neurogenic progenitors (NPs, EGFR^+^PSA-NCAM^−^) and 33.3%±2 neuroblasts (EGFR^−^ PSA-NCAM^+^). **c,** Quantification of OPP incorporation in lineage-traced NSCs and ENBs relative to ENBs. p < 0.0001^****^.

**Extended Data Figure 2.**
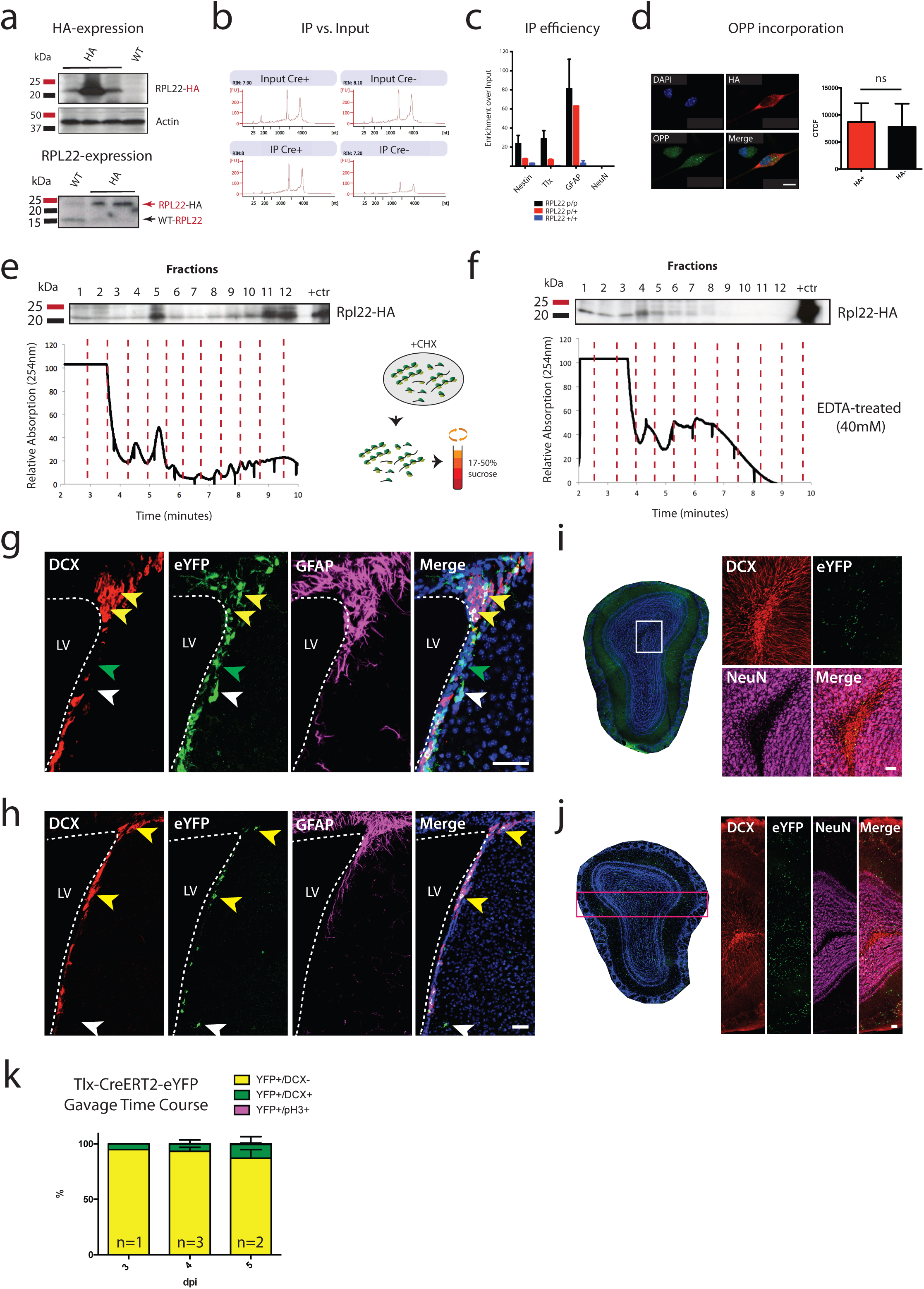
**a,** Top: NSCs isolated from TiCRY mice express HA-tag, replicates show different levels indicating variability based on induction efficiency. Cells of WT mice do not express HA-tag. Bottom: HA-tag is associated to RPL22 since staining shows characteristic 8kDa shift due to HA-tag (23kDa instead of 15kDa). Isolated TiCRY cells entirely replaced WT-RPL22 by RPL22-HA. **b,** Representative Bioanalyzer profiles showing that input RNA levels do not vary between Cre^+^ TiCRY mice (expressing HA-tag) and Cre^−^ TiCRY mice (no HA-tag), however samples from Cre^+^ mice reveal approximately 3-fold more RNA over the background level of Cre^−^ mice after HA-immunoprecipitation. **c,** Enrichment of cell type specific transcripts by qPCR after HA-immunoprecipitation. Efficiency by enrichment of immunoprecipitation-over input samples. Comparison of NSCs from TiCRY mice homozygous for the Rpl22-tag (p/p), heterozygous for the Rpl22-tag (p/+) and WT mice indicates linear relationship between HA-tag expression and immunoprecipitation efficiency. NeuN serves as negative control. Cells isolated from NiCRY mice (Nes-inducible-Cre-Rpl22.HA-eYFP). **d,** OPP incorporation does not vary between HA-tag expressing- and non-expressing cells indicating no major impact on global protein synthesis. NSCs isolated from NiCRY mice. Scale bar: 10µm. n= 30 cells each. **e,** Polysome fractionation using cultured NSCs indicates efficient incorporation of RPL22-HA (antibody against HA-tag) into actively translating polysomes (see fractions 8-12). Total lysate resembles positive control (+ctr). Cells from TiCRY mice. **f,** Disruption of polysomes using EDTA leads to shift of RPL22-HA expression to light fractions indicating specific association to active ribosomes. **g-j,** Immunofluorescence estimating cell type composition based on eYFP expression and characterisitic marker protein expression, see quantification in main figure, all scale bars: 50µm: NSCs using TiCRY mice 4dpi (g), green arrow indicating NSCs, yellow arrows indicating ENB contamination, white arrow showing rare labeled cells outside the SVZ. ENBs using SVZ of DiCRY mice 3dpi (h), yellow arrows pointing at ENBs and white arrow showing rare labeled cells outside the SVZ. LNBs using OB of DiCRY mice 3dpi (i), labeled cells are mostly in the core of OB expressing DCX protein. Neurons using OB of DiCRY mice 4wpi (j), labeled cells throughout the OB mostly coexpressing NeuN protein. **k,** Time course of development of labeled cells in TiCRY mice.

**Extended Data Figure 3.**
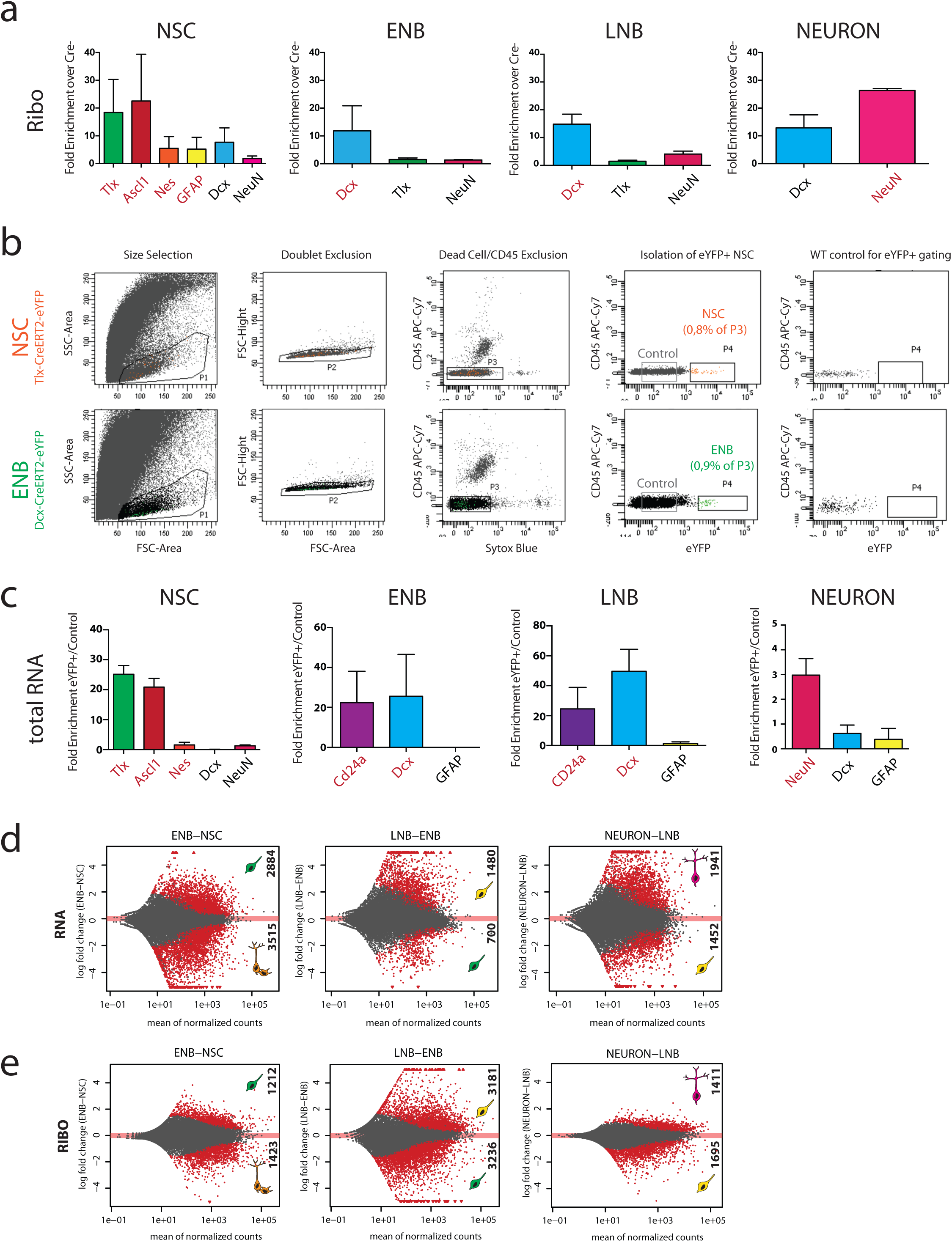
**a,** Enrichment of cell type specific transcripts for NSC, ENB, LNB and Neuron by fold enrichment of HA-IP RNA (RIBO) of Cre^+^ mice over Cre^−^ mice, assessed by qPCR. Genes expected to be strongly enriched after successful immunoprecipitation are labeled in red. **b,** FACS sorting strategy for isolation of NSCs and ENBs based on eYFP expression. eYFP negative cells were collected and used as control samples for enrichment analysis. **c,** Enrichment of cell type specific transcripts by comparison of eYFP^+^ and eYFP^−^ cells by qPCR. Genes expected to be strongly enriched after correct enrichment are labeled in red. All samples were assessed by qPCR before submission for sequencing. **d,** Differential gene expression based on RNAseq. Subsequent populations were compared and significantly changing genes marked in red (FDR: 10%). **e,** Differential gene expression based on RIBOseq without normalization for RIBO-. Subsequent populations were compared and significantly changing genes marked in red (FDR: 10%).

**Extended Data Figure 4.**
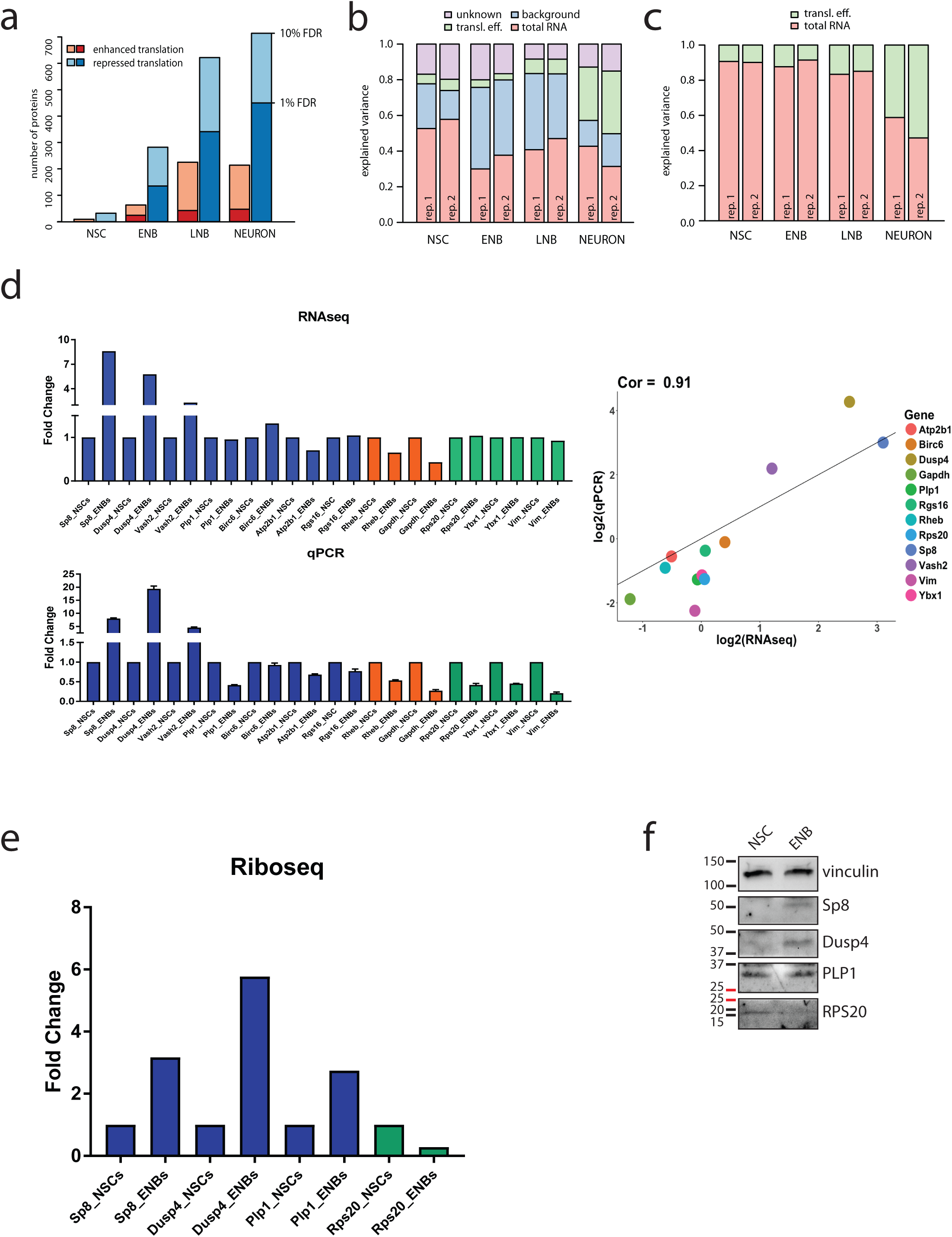
**a,** Summary of the absolute number of genes, which are repressed or enhanced in each population (only protein coding). **b,** Analysis of explained variance scoring the contribution of background (RIBO-), total RNA (RNAseq), translation efficiency (RIBOseq over RNAseq and RIBO-) and unknown residual noise to the RIBOseq data of each population. **c,** As in (b), but comparing only the proportion of variance explained by translation efficiency and RNA. Translation efficiency explains more of the data in neurons compared to earlier developmental stages and NSCs indicating increased importance of translational regulation in neurons. **d**, Correlation in mRNA abundance between the data derived from RNAseq (upper panel) and quantified by qPCR (lower panel) of candidate genes with different translation efficiency upon the NSC-to-ENB transition. The right panel presents a correlation plot for fold changes measured by both approaches. **e**, Fold changes in the RIBOseq fraction of the genes analyzed in (f). **f**, Representative WB image showing relative abundance in freshly sorted NSCs and ENBs of some proteins whose genes change their translation efficiency upon the NSC-to-ENB transition. Vinculin serves as loading control.

**Extended Data Figure 5.**
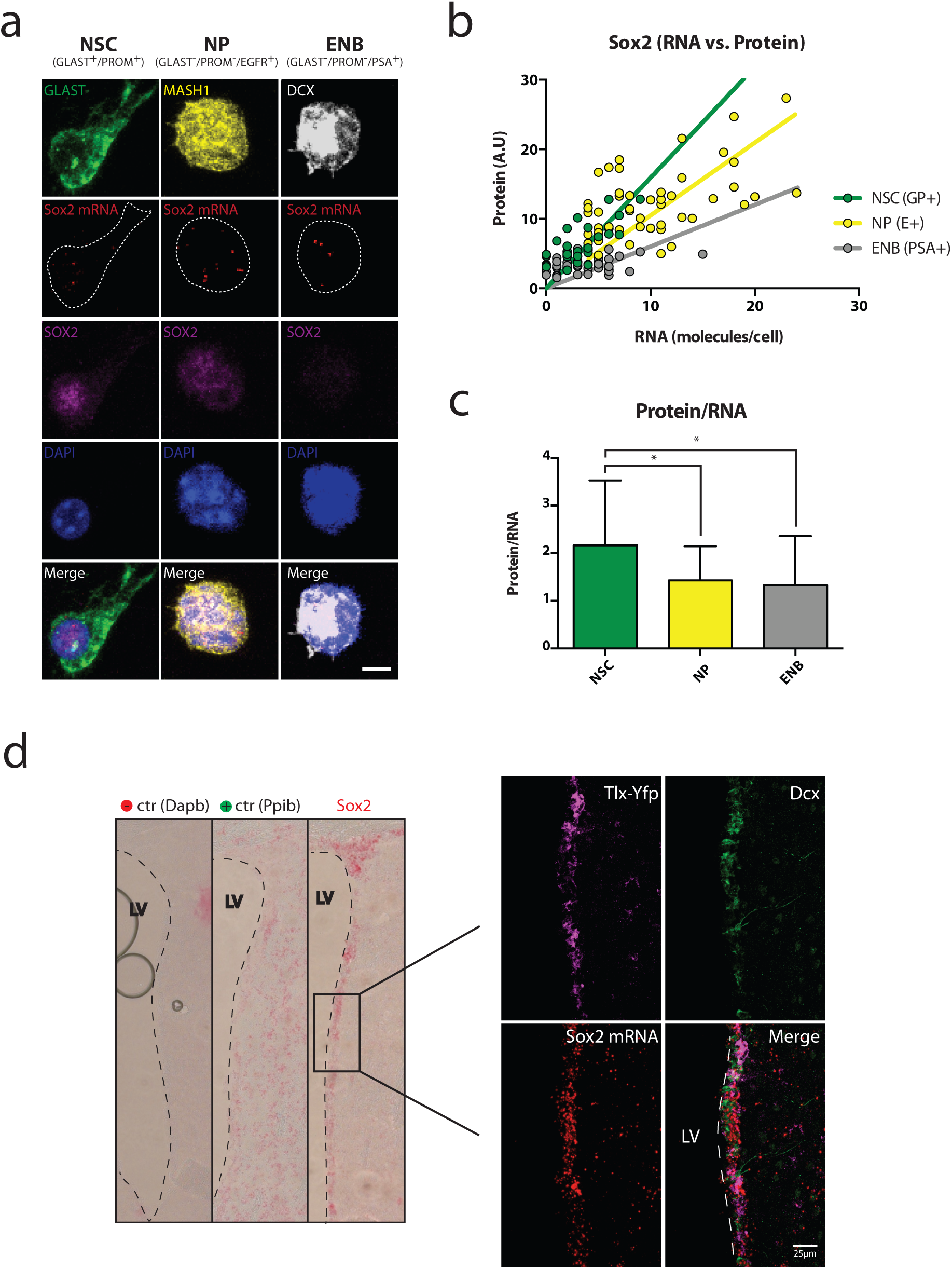
**a,** Representative images for parallel *in situ* hybridization (ISH) and immunocytochemistry of freshly sorted cells of the SVZ. Comparison of NSCs (GLAST^+^PROM^+^), neurogenic progenitors (NPs, GLAST^−^PROM^−^EGFR^+^) and ENBs (GLAST^−^PROM^−^PSA-NCAM^+^). Stained for cell type specific marker proteins and Sox2 protein. ISH for Sox2 mRNA. Scale bar: 5µm. **b,** Quantification of SOX2 protein and RNA expression in NSCs (n=22), NPs (n=49) and ENBs (n=35). Regression line indicates most efficient translation in NSCs. **c,** Ratio of protein expression to RNA abundance indicates higher translation efficiency of Sox2 in NSCs when compared to NPs or ENBs. Significance by students t-test (Mann-Whitney). **d,** Representative image for parallel *in situ* hybridization for Sox2 mRNA in NSCs (marked by Tlx-TOMATO) and ENBs (marked by DCX). Parallel detection of DapB (bacterial transcript) and Ppib (ubiquitous transcriptI) served as a negative- and positive control, respectively. Scale bar: 10µm.

**Extended Data Figure 6.**
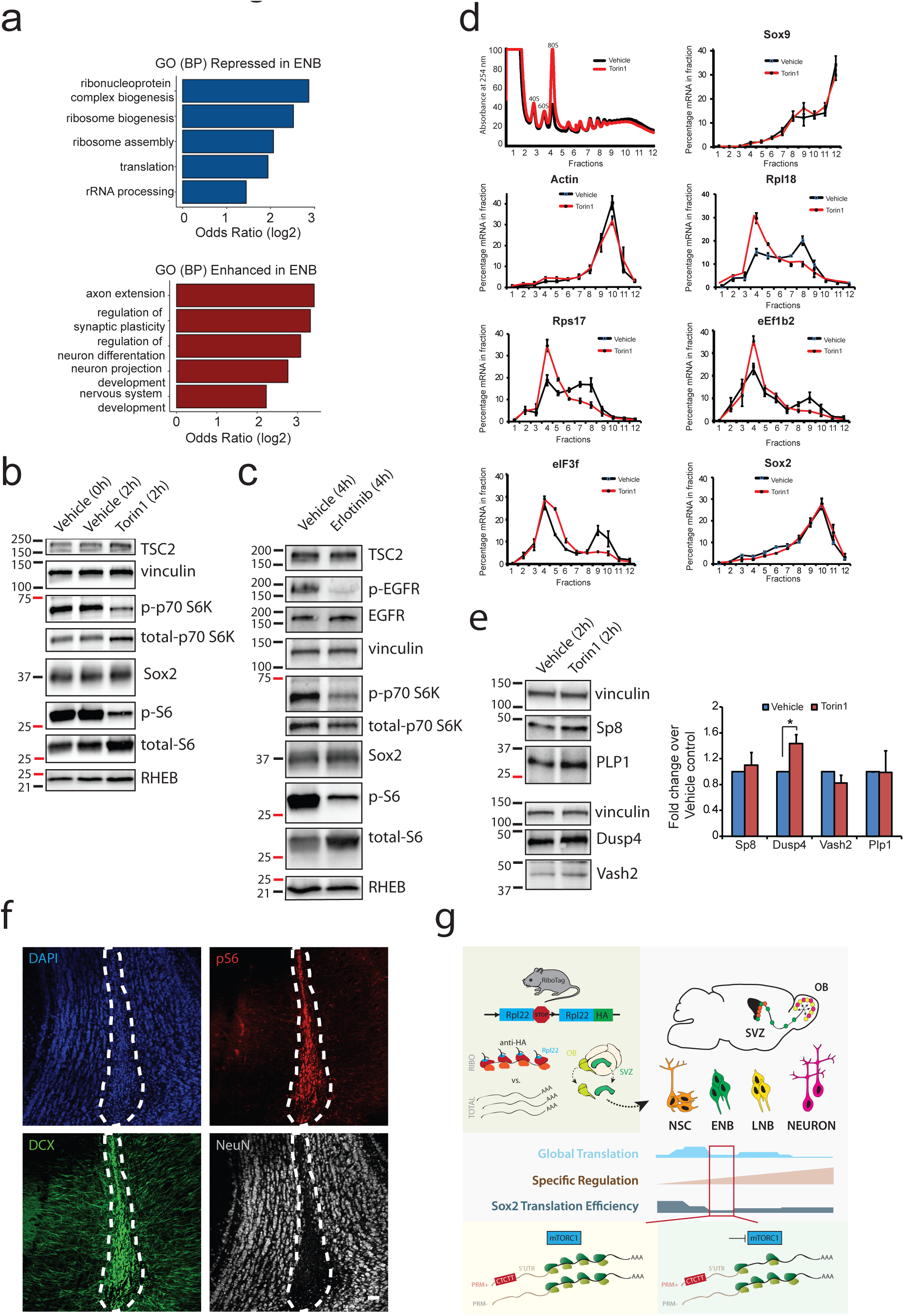
**a,** Gene ontology analysis for genes either repressed (left panel) or upregulated (right panel) at the ENB stage. The repressed genes are mostly related to the protein synthesis machinery, whereas many upregulated genes are involved in development, differentiation and neuronal function. **b, WB of** Torin1-treated primary NSCs showing inhibition of p70 S6 kinase- and S6 phosphorylation, all indicators of mTORC1 activity. mTORC1 regulators TSC2 and RHEB, as well as Sox2, are not affected by 2h incubation with Torin1. Vinculin serves as loading control. **c,** WB of primary NSCs treated for 4h with erlotinib reveals a drop in EGFR phosphorylation accompanied by much lower levels of phospho p70 S6 kinase and phospho S6. TSC2, RHEB and Sox2, similar as for Torin1 treatment, remain at a constant level. **d,** Representative absorption profiles for vehicle- and Torin1-treated NSCs (2h) after sucrose gradient fractionation indicate slight repression in global translation upon torin1 treatment. Representative plots showing the distribution of mRNAs for some genes across the gradient fractions of the polysome profiles are shown. Note a strong shift of PRM+ mRNAs (Rpl18, Rps17, eIF3F, eEF1b2) but not PRM- (Sox9, Actin) to the fractions of light polysomes and non-translating complexes upon Torin1. Sox2 transcript distribution across the polysome fraction was not influenced following 2h Torin treatment **e,** Genes translationally upregulated upon the transition from NSCs to ENBs show slightly higher protein abundance (left panel; WB) but unaffected mRNA levels (right panel; qPCR) following 2h treatment with Torin1 in primary NSCs. **f,** Staining for phosphorylation of ribosomal protein S6 (pS6) in the OB as readout for mTORC1 activity. The core of the OB (marked with dashed line) which contains only LNBs (DCX^+^) is enriched for high pS6, while only few neurons (NeuN^+^DCX^−^) have high levels of pS6. Scale bar: 50µm. **g,** Graphical abstract summarizing the findings of this study.

## REFERENCES

1 Codega, P. et al. Prospective Identification and Purification of Quiescent Adult Neural Stem Cells from Their In Vivo Niche. Neuron 82, 545–559, doi:10.1016/j.neuron.2014.02.039 (2014).

2 Llorens-Bobadilla, E. et al. Single-Cell Transcriptomics Reveals a Population of Dormant Neural Stem Cells that Become Activated upon Brain Injury. Cell Stem Cell 17, 329–340 (2015).

3 Mich, J. K. et al. Prospective identification of functionally distinct stem cells and neurosphere-initiating cells in adult mouse forebrain. eLife 3, 894, doi:10.7554/eLife.02669 (2014).

4 Bayraktar, O. A., Fuentealba, L. C., Alvarez-Buylla, A. & Rowitch, D. H. Astrocyte development and heterogeneity. Cold Spring Harb Perspect Biol 7, a020362, doi:10.1101/cshperspect.a020362 (2015).

5 Ming, G. L. & Song, H. Adult neurogenesis in the mammalian brain: significant answers and significant questions. Neuron 70, 687–702 (2011).

6 Fuentealba, L. C. et al. Embryonic Origin of Postnatal Neural Stem Cells. Cell 161, 1644–1655, doi:10.1016/j.cell.2015.05.041 (2015).

7 Furutachi, S. et al. Slowly dividing neural progenitors are an embryonic origin of adult neural stem cells. Nat Neurosci 18, 657–665, doi:10.1038/nn.3989 (2015).

8 Blanco, S. et al. Stem cell function and stress response are controlled by protein synthesis. Nature 534, 335–340, doi:10.1038/nature18282 (2016).

9 Signer, R. A. J., Magee, J. A., Salic, A. & Morrison, S. J. in Nature Vol. 509 49–54 (2014).

10 Bonaguidi, M. A. et al. In vivo clonal analysis reveals self-renewing and multipotent adult neural stem cell characteristics. Cell 145, 1142–1155 (2011).

11 Rodgers, J. T. et al. mTORC1 controls the adaptive transition of quiescent stem cells from G0 to G(Alert). Nature 510, 393–396 (2014).

12 Liu, J., Xu, Y., Stoleru, D. & Salic, A. in Proc. Natl. Acad. Sci. U.S.A. Vol. 109 413–418 (2012).

13 Jung, H., Yoon, B. C. & Holt, C. E. Axonal mRNA localization and local protein synthesis in nervous system assembly, maintenance and repair. Nature Reviews Neuroscience (2012).

14 Shigeoka, T. et al. in Cell Vol. 166 181–192 (2016).

15 Sanz, E. et al. in Proc. Natl. Acad. Sci. U.S.A. Vol. 106 13939–13944 (2009).

16 Shi, Z. et al. Heterogeneous Ribosomes Preferentially Translate Distinct Subpools of mRNAs Genome-wide. Mol Cell 67, 71–83 e77, doi:10.1016/j.molcel.2017.05.021 (2017).

17 Anger, A. M. et al. Structures of the human and Drosophila 80S ribosome. Nature 497, 80–85, doi:10.1038/nature12104 (2013).

18 Sanz, E. et al. RiboTag analysis of actively translated mRNAs in Sertoli and Leydig cells in vivo. PLoS One 8, e66179, doi:10.1371/journal.pone.0066179 (2013).

19 Cabezas-Wallscheid, N. et al. in Stem Cell 1–16 (Elsevier Inc., 2014).

20 Camp, J. G. et al. Human cerebral organoids recapitulate gene expression programs of fetal neocortex development. Proc Natl Acad Sci U S A 112, 15672–15677, doi:10.1073/pnas.1520760112 (2015).

21 Yamanaka, S. Pluripotency and nuclear reprogramming. Philos.Trans.R.Soc.Lond B Biol.Sci. 363, 2079–2087 (2008).

22 Thoreen, C. C. et al. in Nature Vol. 485 109–113 (2012).

23 Paliouras, G. N. et al. Mammalian Target of Rapamycin Signaling Is a Key Regulator of the Transit-Amplifying Progenitor Pool in the Adult and Aging Forebrain. The Journal of neuroscience: the official journal of the Society for Neuroscience 32, 15012–15026 (2012).

24 Demetriades, C., Doumpas, N. & Teleman, A. A. Regulation of TORC1 in response to amino acid starvation via lysosomal recruitment of TSC2. Cell 156, 786–799, doi:10.1016/j.cell.2014.01.024 (2014).

25 Thoreen, C. C. et al. in J. Biol. Chem. Vol. 284 8023–8032 (2009).

26 Costa, V. et al. mTORC1 Inhibition Corrects Neurodevelopmental and Synaptic Alterations in a Human Stem Cell Model of Tuberous Sclerosis. Cell Rep 15, 86–95, doi:10.1016/j.celrep.2016.02.090 (2016).

27 Sanchez, C. G. et al. Regulation of Ribosome Biogenesis and Protein Synthesis Controls Germline Stem Cell Differentiation. Cell Stem Cell 18, 276–290, doi:10.1016/j.stem.2015.11.004 (2015).

28 Feliciano, D. M., Quon, J. L., Su, T., Taylor, M. M. & Bordey, A. Postnatal neurogenesis generates heterotopias, olfactory micronodules and cortical infiltration following single-cell Tsc1 deletion. Hum Mol Genet 21, 799–810, doi:10.1093/hmg/ddr511 (2012).

29 Li, J. J., Bickel, P. J. & Biggin, M. D. System wide analyses have underestimated protein abundances and the importance of transcription in mammals. PeerJ 2, e270, doi:10.7717/peerj.270 (2014).

30 Schwanhausser, B. et al. Global quantification of mammalian gene expression control. Nature 473, 337–342 (2011).

31 Li, J. J. & Biggin, M. D. in Science Vol. 347 1066–1067 (2015).

32 Gonzalez, C. et al. Ribosome Profiling Reveals a Cell-Type-Specific Translational Landscape in Brain Tumors. The Journal of neuroscience: the official journal of the Society for Neuroscience 34, 10924–10936 (2014).

33 Sahin, E. & DePinho, R. A. Axis of ageing: telomeres, p53 and mitochondria. Nat Rev Mol Cell Biol 13, 397–404, doi:10.1038/nrm3352 (2012).

34 Wei, W. et al. in Cancer Cell Vol. 29 563–573 (2016).

35 Garros-Regulez, L. et al. mTOR inhibition decreases SOX2-SOX9 mediated glioma stem cell activity and temozolomide resistance. Expert Opin Ther Targets 20, 393–405, doi:10.1517/14728222.2016.1151002 (2016).

36 Pelletier, J., Graff, J., Ruggero, D. & Sonenberg, N. Targeting the eIF4F translation initiation complex: a critical nexus for cancer development. Cancer Res 75, 250–263, doi:10.1158/0008-5472.CAN-14-2789 (2015).

37 Hsieh, A. C. et al. The translational landscape of mTOR signalling steers cancer initiation and metastasis. Nature 485, 55–61, doi:10.1038/nature10912 (2012).

38 Shatsky, I. N., Dmitriev, S. E., Andreev, D. E. & Terenin, I. M. Transcriptome-wide studies uncover the diversity of modes of mRNA recruitment to eukaryotic ribosomes. Crit Rev Biochem Mol Biol 49, 164–177, doi:10.3109/10409238.2014.887051 (2014).

39 Meyer, K. D. & Jaffrey, S. R. Rethinking m6A Readers, Writers, and Erasers. Annu Rev Cell Dev Biol, doi:10.1146/annurev-cellbio-100616-060758 (2017).

40 Liu, H. K. et al. The nuclear receptor tailless is required for neurogenesis in the adult subventricular zone. Genes Dev. 22, 2473–2478 (2008).

41 Werner, L. et al. Involvement of doublecortin-expressing cells in the arcuate nucleus in body weight regulation. Endocrinology 153, 2655–2664 (2012).

42 Retz, K. C. & Steele, W. J. in Life Sci. Vol. 27 2601–2604 (1980).

43 Mirzadeh, Z., Doetsch, F., Sawamoto, K., Wichterle, H. & Alvarez-Buylla, A. in JoVE (2010).

44 Picelli, S. et al. in Nat Protoc Vol. 9 171–181 (2014).

45 Dinkel, C., Moody, M., Traynor-Kaplan, A. & Schultz, C. Membrane-Permeant 3-OH-Phosphorylated Phosphoinositide Derivatives. Angew Chem Int Ed Engl 40, 3004–3008, doi:10.1002/1521-3773(20010817)40:16<3004::AID-ANIE3004>3.0.CO;2-O (2001).

46 Lou, W. P., Baser, A., Klussmann, S. & Martin-Villalba, A. In vivo interrogation of central nervous system translatome by polyribosome fractionation. J.Vis.Exp. (2014).

47 Dobin, A. et al. in Bioinformatics Vol. 29 15–21 (2013).

48 Li, B. & Dewey, C. N. in BMC Bioinformatics Vol. 12 323 (2011).

49 Love, M. I., Huber, W. & Anders, S. in Genome Biol. Vol. 15 550 (2014).

50 Ritchie, M. E. et al. in Nucleic Acids Research Vol. 43 e47 (2015).

